# The mechanism of cysteine sensing by LRRC58

**DOI:** 10.64898/2026.11.03.763985

**Authors:** Emma C. Fink, Hilina B. Woldemichael, Nikhit Kambdur, Muneeb A. Sharif, Talya S. Levitz, Bindu Srinivasu, Marie Blickling, Bingsen Zhang, Jiuchun Zhang, Yaoyu Wang, Filipe Cabreiro, Thomas E. Wales, Eric S. Fischer, Edward T. Chouchani

## Abstract

Cysteine is essential for life but toxic in excess. Cells and tissues are exposed to elevated cysteine from endogenous sources and the environment yet maintain low intracellular concentrations to sustain life. Here we identify a mechanism for sensing cysteine to control its abundance and prevent toxicity and premature death. Using cryo-electron microscopy we determine structures of the CRL5^LRRC58^ E3 ligase alone and in complex with the cysteine-catabolizing enzyme CDO1. We show that CDO1 interacts with LRRC58 in the vicinity of a zinc-coordinating cysteine motif, within which Cys296 becomes covalently modified upon cysteine exposure. Covalent modification of Cys296 and zinc ejection from this motif drive large structural rearrangements of LRRC58 relayed through Cys319, which destabilize CDO1 binding and position LRRC58 Lys111 for autoubiquitylation. This reorientation of LRRC58 permits CDO1 accumulation, thereby lowering intracellular cysteine. Mutation of Cys319 abrogates inhibitory rearrangement of LRRC58 upon cysteine modification. This renders the ligase constitutively active and prevents CDO1 stabilization, thereby driving systemic cysteine accumulation and premature death in animals. These findings establish the molecular basis of homeostatic cysteine sensing by LRRC58 and demonstrate its essential role in maintaining cysteine concentrations compatible with healthy life.

---

The sulfur-containing amino acid cysteine plays a vital role in numerous biological processes and is essential for life. Cysteine is produced endogenously from transsulfuration and protein breakdown or is obtained directly from dietary sources. Cysteine is a necessary precursor for protein synthesis and various essential biomolecules including glutathione (GSH), coenzyme A, and inorganic sulfur-containing compounds. While essential, cysteine’s cellular abundance must be strictly regulated as its accumulation is toxic, a phenomenon observed widely across evolution^1–7^. Elevated intracellular cysteine drives profound mitochondrial dysfunction and promotes iron-dependent cytotoxicity^1,6,8^. Excess cysteine is damaging to animal tissues^9–12^ and is associated with numerous chronic diseases in humans. In particular, elevated cysteine is associated with rheumatoid arthritis^13^, neurodegenerative diseases^14^, cardiometabolic disorders^15,16^, and increased biological aging and mortality^17^.

Cysteine’s dual nature as both an essential metabolite and one that becomes toxic in excess necessitates mechanism(s) that sense and control its abundance. It has been appreciated for decades that healthy cells and tissues maintain low concentrations of cysteine^18,19^ despite frequent fluctuations in endogenous cysteine production and exposure to cysteine from the diet^19^. While these longstanding observations support the existence of a putative cysteine sensor, a mechanistic basis for cysteine sensing has remained elusive.

As cellular cysteine is derived from multiple sources, we reasoned that an efficient mechanism for maintaining low cysteine concentrations would act through regulation of its catabolism. The enzyme cysteine dioxygenase 1 (CDO1) is a major catabolic route for cysteine in cells. Seminal studies by Martha Stipanuk and colleagues demonstrated that CDO1 protein abundance is sensitive to dietary cysteine intake. For example, CDO1 protein and activity in the liver are barely detectable on sulfur-depleted diets, but increase rapidly upon dietary intake of cysteine or cysteine precursors^20,21^. Remarkably, these cysteine-driven changes in CDO1 protein abundance are post-translational and depend on the ubiquitin-proteasome system^22^. Together, these findings suggest that a molecular rheostat exists to regulate intracellular cysteine levels through degradation of CDO1.

We discovered that LRRC58 is the substrate receptor of an E3 ubiquitin ligase that targets CDO1 for proteasomal degradation^23^; a finding subsequently corroborated by independent studies^24,25^. In cells, LRRC58 assembles with CUL2 or CUL5, RBX1 or RBX2, ELOB, and ELOC (CRL^LRRC58^) to ubiquitylate CDO1 and mediate its subsequent degradation by the proteasome. This process is active when cysteine is scarce and inhibited when cysteine is supplied to cells^23–25^. These studies established the machinery responsible for CDO1 turnover but left unresolved the central question of how cysteine is sensed: does CRL^LRRC58^ detect cysteine directly, or does it respond to a cysteine-derived signal or an upstream intermediary? Moreover, whether this complex maintains cysteine homeostasis and protects cells from cysteine toxicity also remained unknown. Here, using structural, biophysical, and biochemical approaches, we identify LRRC58 itself as a direct cysteine sensor and reveal how cysteine recognition controls CDO1 degradation. We further demonstrate that this sensing mechanism is essential to restrain intracellular cysteine accumulation and sustain organismal viability, establishing both the molecular basis and physiological importance of this cysteine homeostatic circuit.

## RESULTS

### Cyst(e)ine disrupts the interaction between the CRL5^LRRC58^ E3 ligase complex and CDO1

In cells, LRRC58-dependent degradation of CDO1 is potentiated when cysteine is restricted and strongly antagonized when cysteine accumulates. This indicates that cysteine, or a metabolic product of cysteine, somehow antagonizes the function of the CRL^LRRC58^ E3 ligase complex. This regulation could be direct or mediated indirectly through some undefined intermediary. Applying the principle of parsimony^26^, our favored hypothesis was that cysteine acts directly on CRL^LRRC58^ to inhibit its capacity to ubiquitylate CDO1. To test this, we reconstituted the full CRL5^LRRC58^ complex (LRRC58, ELOB, ELOC, CUL5, RBX2) from recombinantly expressed proteins. Upon co-incubation, CRL5^LRRC58^ and recombinant CDO1 formed a ∼203 kDa complex **(fig. S1A)**, and spontaneous interaction between CRL5^LRRC58^ and CDO1 was observed by co-immunoprecipitation (co-IP) of CRL5^LRRC58^ **(Fig. 1A,C)**. Addition of cysteine to the reconstituted complex disrupted the CRL5^LRRC58^-CDO1 interaction in a concentration-dependent manner across the physiologic range of cysteine, with maximal effect at 500 μM cysteine **(Fig. 1A,B)**. Cysteine specifically diminished the CDO1-LRRC58 interaction and had no observable effect on the other contacts within CRL5^LRRC58^ **(Fig. 1A)**. Intracellular cysteine exists in equilibrium with its oxidized form cystine, so we examined the effects of cystine on CRL5^LRRC58^-CDO1. Like cysteine, cystine disrupted the interaction between CDO1 and CRL5^LRRC58^ **(Fig. 1C,D)**.

**Figure 1.**
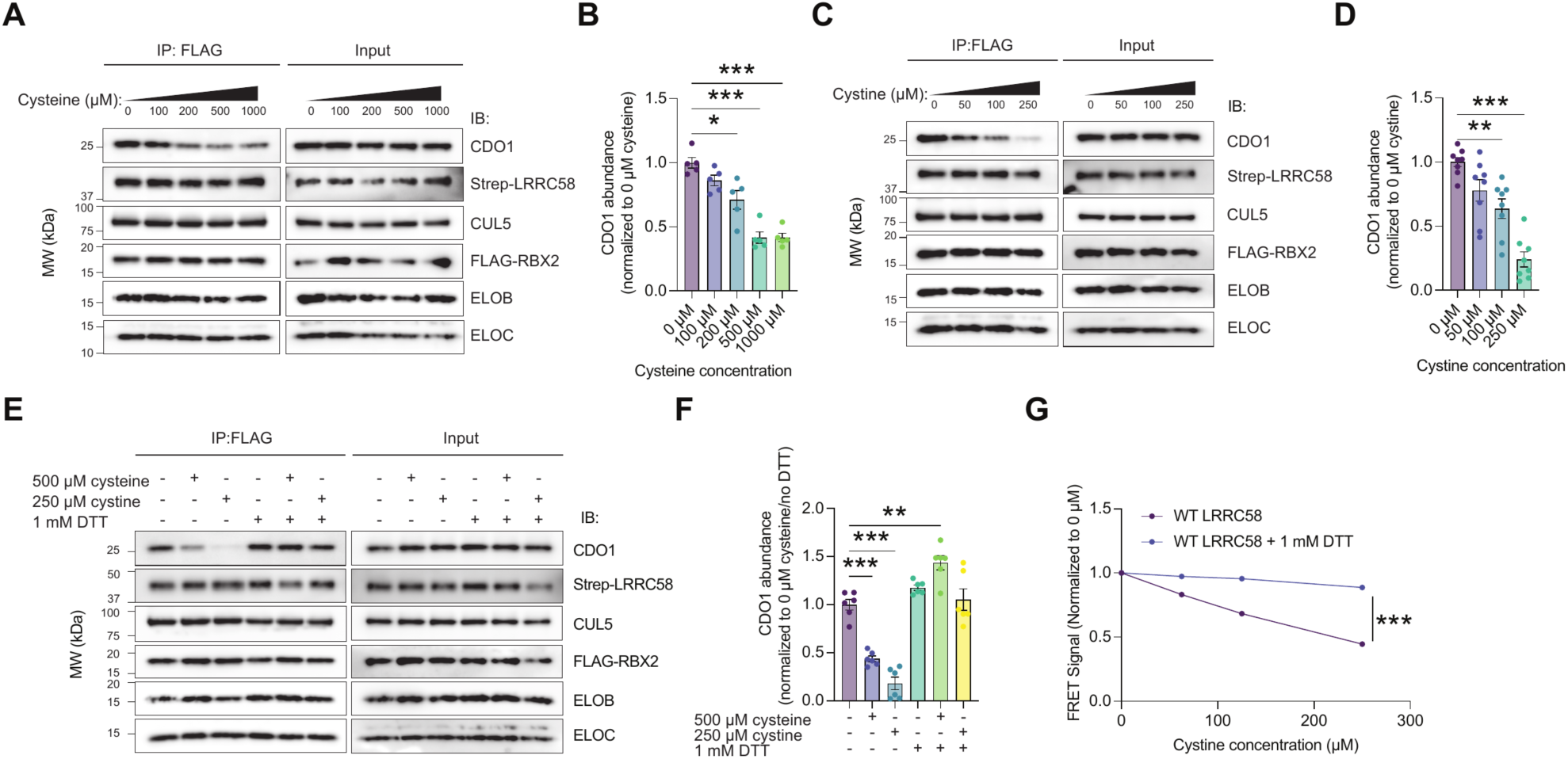
Cysteine and cystine directly disrupt the CRL5^LRRC58^-CDO1 interaction through a reversible thiol-covalent mechanism. **(A)** Representative FLAG IP immunoblot analysis of interaction between recombinantly expressed and purified preassembled CRL5^LRRC58^ (CUL5-FLAG∼RBX2-Strep∼LRRC58-ELOB-ELOC) and recombinantly expressed and purified Strep∼CDO1 following 1 h preincubation in buffer with increasing concentrations of L-cysteine. **(B)** Densitometric quantification of CDO1 abundance from the immunoblots in (**A**) as a function of increasing cysteine concentration. *n* = 5 replicates from 3 independent experiments. **(C)** Representative FLAG IP immunoblot analysis of binding between recombinant preassembled CRL5^LRRC58^ and recombinant Strep∼CDO1 following 1 h preincubation in buffer with increasing concentrations of L-cystine. **(D)** Densitometric quantification of CDO1 abundance from the immunoblots in (**C**) as a function of increasing cystine concentration in the reaction buffer. *n* = 8 replicates pooled from 4 independent experiments. **(E)** Representative FLAG IP immunoblot analysis of interaction between recombinant preassembled CRL5^LRRC58^ and recombinant Strep∼CDO1 following 1 h preincubation in buffer, cysteine (500 µM) or cystine (250 µM), ± 1 mM DTT. **(F)** Densitometric quantification of CDO1 abundance from the immunoblots in (**E**). *n* = 6 replicates from 3 independent experiments. **(G)** Time-resolved Förster resonance energy transfer (TR-FRET) signal between eGFP-LRRC58-ELOB-ELOC and CDO1 across increasing concentrations of L-cystine ± 1 mM DTT. *n* = 3, representative of 2 independent experiments. Statistics: Data are mean ± SEM. Asterisks denote * *P* < 0.05, ** *P* < 0.01, *** *P* < 0.001. Brown-Forsythe and Welch ANOVA with Dunnett’s T3 correction for panels (**B**), (**D**), (**F**). Two-way ANOVA for (**G**), *P* <0.0001 for cystine dose, DTT, and their interaction. Dunnett’s test: *P* < 0.0001 for each-DTT dose compared to 0 µM. Sidak’s test: *P* < 0.0001, cystine vs. cystine + DTT.

Both cysteine and cystine participate in thiol redox chemistry and can promote reversible covalent modification of protein cysteine residues^27,28^. These reversible thiolation modifications can form stably or act as a transient intermediate before collapsing to a protein disulfide^27^. Covalent modifications of this nature require an appropriately oriented redox-active cysteine on the target protein and are reversed by thiol reductants. Notably, the thiol reductant dithiothreitol (DTT) completely abrogated the effects of both cysteine and cystine on the CRL5^LRRC58^-CDO1 interaction **(Fig. 1E,F)**. These data suggest that cysteine and cystine could disrupt the interaction between CRL5^LRRC58^ and CDO1 through reversible covalent modification of one or several protein cysteines on the E3 ligase complex or on CDO1. We next tested whether these effects were specific to cyst(e)ine. We profiled major cellular metabolic products of cysteine and related redox-active metabolites. None of these metabolites significantly disrupted the interaction between CRL5^LRRC58^ and CDO1 **(fig. S1B,C)**.

We corroborated the direct effect of cystine on the LRRC58-CDO1 interaction using a time-resolved Förster resonance energy transfer (TR-FRET) binding assay. We observed a dissociation constant (K_d_) of ∼ 300 ± 12 nM for the interaction between eGFP-tagged LRRC58-ELOB-ELOC (LRRC58_eGFP_) and terbium-chelate-labeled CDO1 (CDO1_Tb_) **(fig. S1D)**.

Application of cystine resulted in concentration-dependent disruption of the LRRC58_eGFP_-CDO1_Tb_ complex across the physiologic range **(Fig. 1G)**. Again, disruption of the complex by cystine was prevented by addition of the thiol reductant DTT **(Fig. 1G)**. Taken together, our findings indicated that cysteine and cystine directly disrupt the interaction between the CRL5^LRRC58^ E3 ligase complex and CDO1 through a thiol redox mechanism.

### Structure of CRL5^LRRC58^-CDO1 defines a cysteine-rich motif proximal to LRRC58’s substrate recognition interface

To understand how cyst(e)ine could selectively disrupt the interaction between CDO1 and CRL5^LRRC58^, we determined the cryo-electron microscopy (cryo-EM) structure of the CRL5^LRRC58^-CDO1 complex **(fig. S2)**. Several rounds of classification and local refinements led to cryo-EM maps refined to resolutions from 2.92 to 3.13 Å sufficient for high-confidence model building (**Fig. 2A** & **S2** & **table S1)**. The CRL5^LRRC58^-CDO1 complex adopts a canonical cullin-RING ligase organization **(Fig. 2A)**, with the ELOB/ELOC adaptor bridging CUL5 and the substrate receptor LRRC58 through interactions with both proteins. This arrangement positions CDO1 at the N-terminal end of the CUL5 scaffold, opposite to the catalytic RBX2 RING domain, and poised for ubiquitylation **(Fig. 2A)**. In support of this, during preparation of this manuscript, a cryo-EM structure of a covalently trapped mimic of a CRL^LRRC58^-CDO1 ubiquitin-transfer intermediate (PDB 9T7V)^25^ was reported. While the structures agree closely, our structure captures the native substrate-engaged state of the ligase, whereas the trapped complex is held in a catalytically poised configuration.

**Figure 2.**
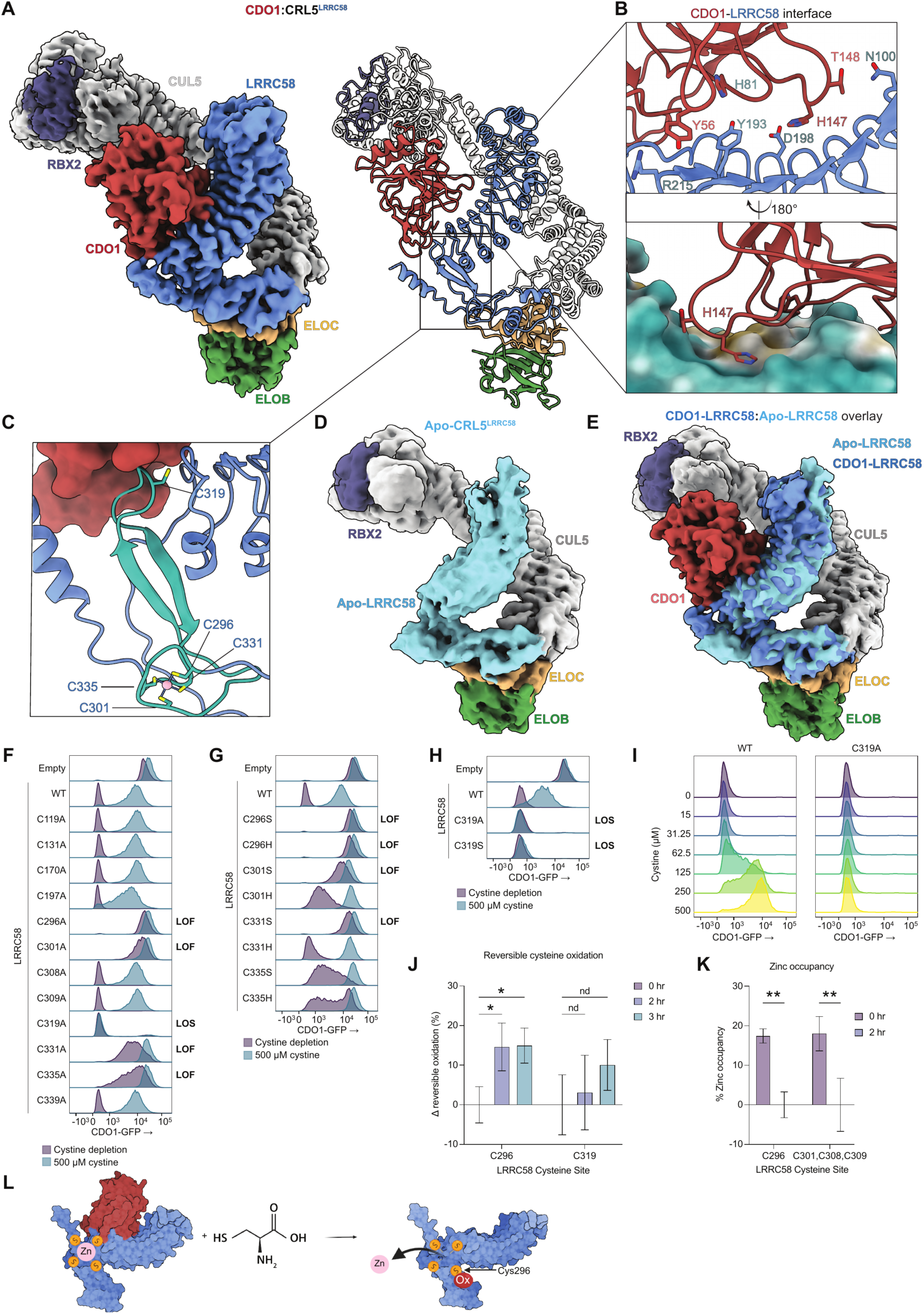
LRRC58 zinc-coordinating cysteine cluster adjacent to the CDO1 interface is oxidized and loses zinc upon cyst(e)ine exposure. **(A)** Cryo-EM map (left) and atomic model shown as cartoon representation (right) of the CDO1-CRL5^LRRC58^ complex. Subunits are colored and labeled: CUL5 (grey), RBX2 (dark purple), Elongin B (ELOB) [green], Elongin C (ELOC) [light orange], LRRC58 (blue), and CDO1 (red) [Map shown at a contour level of 12 σ; overall resolution 2.92-3.13 Å — see **fig. S2**]. **(B)** Close-up of the CDO1–LRRC58 binding interface. CDO1 (red) engages the concave face of the LRRC58 (blue) leucine-rich repeat solenoid. Top, zoomed view of the boxed interface with key contact residues shown as sticks and labeled (CDO1, red; LRRC58, blue). Bottom, H147 of CDO1 with electrostatic surface of LRRC58. **(C)** Cryo-EM structure of CRL5^LRRC58^-CDO1 with LRRC58 zinc-binding cysteine cluster (Cys296, Cys301, Cys331, Cys335) and Cys319 indicated. The region containing these residues is highlighted in teal with CDO1 shown as a surface representation (red). **(D)** Cryo-EM map of Apo-CRL5^LRRC58^. **(E)** Alignment of Apo structure and CDO1-LRRC58, with LRRC58 in Apo map colored light blue and LRRC58 in CDO1-LRRC58 map colored blue. **(F-H)** CDO1-eGFP fluorescence normalized to mode in the mCherry^+^ population of LRRC58 KO Hep G2 cells expressing the WT CDO1 post-translational stability reporter and reconstituted with the indicated single-cysteine LRRC58 point mutants (or empty vector/WT), after 24 h in cystine-depleted (purple) or 500 µM cystine (teal) medium. LOF, loss-of-function; LOS, loss-of-sensing. (**F**) All LRRC58 cysteines substituted with alanine. (**G**) Ser/His substitutions at the LOF cysteines identified in (**F**). (**H**) C319A and C319S. Data are one representative replicate of ≥3 independent experiments. **(I)** CDO1-eGFP fluorescence normalized to mode in mCherry^+^ population of cell lines generated as in (**F-H**) and exposed to media with the indicated dose of cystine for 24 hr. Data are one representative replicate of 3 independent experiments. **(J)** Change in percent reversible oxidation at the indicated LRRC58 cysteines in FLAG-LRRC58 immunoprecipitated from Hep G2 cells maintained in cystine depletion overnight, then treated with 200 µM cystine for 0, 2, or 3 h in the presence of MG132. Data are normalized to 0 h by subtraction; *n* = 8-9 cell replicates. (K) Percent zinc-occupancy of zinc-coordinating cysteines in FLAG-LRRC58 immunoprecipitated from Hep G2 cells maintained in cystine depletion overnight then treated with 200 µM cystine for 0 or 2 hr in the presence of MG132. *n* = 5 cell replicates per time point. **(L)** Model of zinc removal from 4-Cys motif in LRRC58 following cysteine exposure and oxidation at Cys296. Statistics: Data are mean ± SEM unless otherwise noted. (**J**) Kruskal–Wallis test with Dunn’s multiple comparisons of the 2 h and 3 h timepoints to 0 h. False discovery rate controlled at 5% by two-stage step-up procedure of Benjamini, Krieger and Yekutieli. * *q* < 0.05. nd, no discovery. (**K**) Two-tailed Welch’s *t*-test on unbounded values (Holm-Šídák correction). \*\**P* < 0.01.

The leucine-rich repeat (LRR) domain of LRRC58 forms a curved solenoid, with a concave inner surface lined by a parallel β-sheet and a convex outer surface composed of helices and loops. The C-terminal region of LRRC58 contains a BC-box motif that mediates interaction with ELOC, linking the substrate receptor and the CUL5 scaffold. CDO1 binds to the concave face of the LRRC58 LRR domain. On the LRRC58 side, the interface is mediated by residues from the concave β-sheet surface and a C-terminal segment **(Fig. 2B)**. On CDO1, the interface involves residues from multiple surface loops and β-strands stabilized by a combination of hydrogen bonds, salt bridges, and hydrophobic contacts, including His147 that forms an amide-π interaction with the backbone amide of LRRC58 Gly175 **(Fig. 2B)**. This provides a structural basis for our previous observation that CDO1 His147 is required for destruction of CDO1 in cells^23^. Indeed, the interaction between LRRC58_eGFP_ and CDO1_Tb_ was completely lost upon mutation of CDO1 His147 to alanine **(fig. S3A)**.

To validate the observed CDO1–LRRC58 interface in solution and probe the local conformational dynamics of the complex, we employed hydrogen–deuterium exchange mass spectrometry (HDX-MS). HDX-MS measures the exchange of backbone amide hydrogens with deuterium. Increased deuterium exchange (deprotection) indicates greater solvent exposure or conformational flexibility through the loss or weakening of backbone hydrogen-bonding. Conversely, decreased exchange (protection) indicates burial, hydrogen-bond stabilization, or altered backbone environment due to a protein–protein interface. Comparison of CRL5^LRRC58^-CDO1 in complex with the individual proteins revealed increased protection on both LRRC58 and CDO1 at the interface identified by cryo-EM, confirming the LRRC58-CDO1 interface in solution **(fig. S3B-G)**.

Given that cysteine disrupted LRRC58 interaction with CDO1 through a thiol redox mechanism **(Fig. 1)**, a notable feature of the cryo-EM structure is the presence of a cysteine-rich region proximal to the LRRC58-CDO1 interface. This region includes a zinc coordination site comprising Cys296, Cys301, Cys331, and Cys335 located at the hinge between the BC-box and the LRR domain **(Fig. 2C)**. This 4Cys zinc coordination staples the base of a loop that projects ∼28 Å from the metal toward CDO1 **(Fig. 2C)**. Notably, Cys319 sits at the apex of this loop at the periphery of the CDO1 interface. The zinc-coordinating cysteines Cys296 and Cys301 precede this loop, while Cys331 and Cys335 are at the end of the loop forming a kink leading into a short C-terminal section clearly defined in our structure. We hypothesized that this cysteine-rich region of LRRC58 may be a target of cysteine action, as it forms a metastable arrangement poised for conformational change that could in turn alter the interaction with CDO1.

To better understand structural features of the ligase that depend on CDO1 binding, we also determined the Apo structure of CRL5^LRRC58^ in the absence of CDO1 by cryo-EM **(Fig. 2D & S4, table S1;** overall resolution 3.92 Å). While the overall resolution of the Apo structure was reduced by the increase in flexibility of the LRRC58 module, we were still able to rigid body fit the CRL5^LRRC58^ model with high confidence. Aligned on the CUL5-ELOB-ELOC scaffold, the two structures are essentially superimposable across the complex, including the cysteine-rich domain of LRRC58 **(Fig. 2E)**, suggesting that Apo CRL5^LRRC58^ is competent to bind CDO1 without prior structural rearrangements, consistent with surveillance of CDO1 levels at baseline.

By identifying cysteine-rich regions proximal to the CDO1-LRRC58 interface, our structures provide clues as to how cysteine could act directly to regulate CDO1 degradation through a covalent mechanism. We next sought to reconcile these structures with our biochemical observations that cysteine and cystine disrupt the CRL5^LRRC58^-CDO1 interaction through a DTT-reversible mechanism **(Fig. 1E-G)** and determine how this complex could serve as a molecular sensor of cysteine abundance.

### Cysteine drives modification of LRRC58 Cys296 and zinc ejection

Cryo-EM defined the surface residues of LRRC58 that mediate recognition and binding to CDO1 **(Fig. 2B)**. Notably, this interface is adjacent to a region in LRRC58 that appears stabilized by a 4Cys zinc-coordination site, which is proximal to three additional solvent-exposed cysteines **(Fig. 2C)**. Thiol reductants abrogated the cysteine-mediated dissociation of CDO1 from LRRC58. Based on this, we hypothesized that one or more cysteine residues on LRRC58 or CDO1 could be targets for cysteine-driven redox modifications that mediate cysteine sensing by inhibiting LRRC58-mediated ubiquitylation of CDO1.

To test this hypothesis, we first determined whether any cysteine residues on LRRC58 or CDO1 are required for LRRC58-mediated CDO1 degradation (essential for function), or inhibition of this LRRC58 activity by cysteine (essential for sensing). We used a fluorescent reporter of CDO1 post-translational stability that enables real-time monitoring in live cells **(fig. S5A,B)**. In this context we manipulated intracellular cysteine concentrations through titration of cysteine equivalents in culture media supplied as cystine, which is taken up by plasma membrane transporters and reduced to cysteine intracellularly^6,29^. As shown previously^23^, media cystine depletion rapidly decreased CDO1 post-translational stability in an LRRC58-dependent manner, whereas elevation of extracellular cysteine or cystine inhibited LRRC58 and stabilized CDO1 **(fig. S5C)**. We next generated derivative reporters in which each CDO1 cysteine residue was individually mutated to alanine. All CDO1 mutants were indistinguishable from WT, demonstrating that no individual CDO1 cysteine residue is required for LRRC58-mediated degradation or cysteine sensing **(fig. S5D)**.

To assess the importance of LRRC58 cysteine residues, we introduced the WT CDO1 reporter into LRRC58 KO cells complemented with each of 12 cysteine-to-alanine LRRC58 variants. While most LRRC58 cysteine mutants were indistinguishable from WT, mutation of any of the zinc-coordinating cysteines (Cys296, Cys301, Cys331, and Cys335) to alanine resulted in loss of LRRC58-mediated destruction of CDO1 **(Fig. 2F & S5E)**. Interestingly, substitution of Cys301, Cys331, or Cys335 with histidine, an alternative zinc-coordinating residue, maintained both CDO1 degradation and cysteine sensing **(Fig. 2G)**. Therefore, a cysteine side chain at these positions is not required for LRRC58 function or cysteine sensing, but zinc coordination by this cluster appears to be required for LRRC58-mediated destruction of CDO1. In contrast, Cys296 was intolerant to histidine substitution, suggesting that this position uniquely requires a cysteine side chain for LRRC58 function **(Fig. 2G)**.

Cys319 exhibited a distinct phenotype compared to all other LRRC58 cysteines. Replacement of Cys319 with alanine or serine preserved capacity for CDO1 degradation but completely abolished cysteine sensing **(Fig. 2F,H)**. In cells expressing LRRC58 C319A, CDO1 degradation was invariant across a broad range of extracellular cystine concentrations and exposure durations, indicating a complete loss of sensing rather than a shift in sensitivity or kinetic response to intracellular cysteine **(Fig. 2I & S5F)**. Thus, C319A uncouples LRRC58 degradation activity from its regulation by cysteine, highlighting this LRRC58 residue as a principal determinant of cysteine sensing.

Having identified Cys296 and Cys319 as the only cysteine residues required for LRRC58 function or cysteine sensing, we reasoned that these sites might undergo reversible covalent modification, consistent with the ability of thiol reductants to prevent cysteine-induced dissociation of CDO1 from LRRC58. To test this, we exposed Hep G2 cells to cystine at a concentration sufficient to inhibit LRRC58-dependent CDO1 degradation and quantified reversible covalent modifications on LRRC58 Cys296 and Cys319 through differential alkylation and mass spectrometry (MS) **(fig. S5G,H)**. Cys296 showed a significant increase in reversible covalent modification after both 2 h and 3 h of cystine repletion, while there was no significant change in the modification state of Cys319 **(Fig. 2J)**.

Our cryo-EM structure showed that Cys296 coordinates zinc along with Cys301, Cys331, and Cys335. We therefore hypothesized that cysteine-driven covalent modification of Cys296 would drive collapse of the zinc coordination site and ejection of zinc. The notion that the LRRC58 zinc coordination site is important for CDO1 engagement was supported by the observation that zinc chelation disrupted the interaction between reconstituted CRL5^LRRC58^ and CDO1 **(fig. S5I)**. We used a quantitative MS approach (adapted from ref. ^30^) to measure zinc occupancy at the LRRC58 zinc coordination site in cells **(fig. S5J)**. Both Cys296 and Cys301-containing tryptic peptides could be readily detected and assessed for zinc occupancy. Notably, we found that under low cystine conditions an equivalent fraction of both Cys296 and Cys301 was zinc-occupied **(Fig. 2K)**. Upon application of cystine at a concentration sufficient to inhibit LRRC58-mediated degradation of CDO1, we observed a decrease of zinc occupancy at Cys296 and Cys301 to undetectable levels **(Fig. 2K)**. Together, our data indicate that cyst(e)ine drives covalent modification of Cys296 and loss of zinc coordination at this LRRC58 4Cys site **(Fig. 2L)**.

### Cyst(e)ine modification collapses the zinc site and rotates LRRC58 relative to the CUL5 scaffold

To determine how cyst(e)ine modification remodels LRRC58 and disrupts its activity towards CDO1, we paired cryo-EM with HDX-MS on the cystine-modified CRL5^LRRC58^ complex. Cystine was used in place of cysteine for these experiments as it disrupts the LRRC58-CDO1 interaction at a lower concentration **(Fig. 1B,D)**. We incubated CRL5^LRRC58^ with cystine for 2 h immediately prior to grid preparation and collected cryo-EM data. 2D classification and extensive rounds of 3D classification produced maps with clearly defined high-resolution density for the Cullin and ELOB/ELOC module **(Fig. 3A & S6)**. Heightened mobility of LRRC58 relative to the Cullin-ELOB/ELOC precluded high-resolution reconstruction and limited model building to rigid body fitting of the substrate receptor itself **(Fig. 3A,B & S6, table S1 & movie S1)**, but this flexibility was itself informative. Although the resolution was insufficient to confidently resolve the C-terminal cysteine-rich region, an overlay of the maps and fit of the CDO1-bound model into the cystine map shows that the 4Cys zinc-coordination motif is absent from the position observed in the CDO1-bound state **(Fig. 3B)**. In the cystine map, density for residues 293 to 353 is absent, and neither the motif nor the zinc can be modeled **(Fig. 3B)**. By comparison, the Apo complex lacking CDO1 retains the 4Cys motif and its bound zinc, so loss of this density in the cystine map is a result of cystine action rather than the absence of substrate or a limitation of resolution.

**Figure 3.**
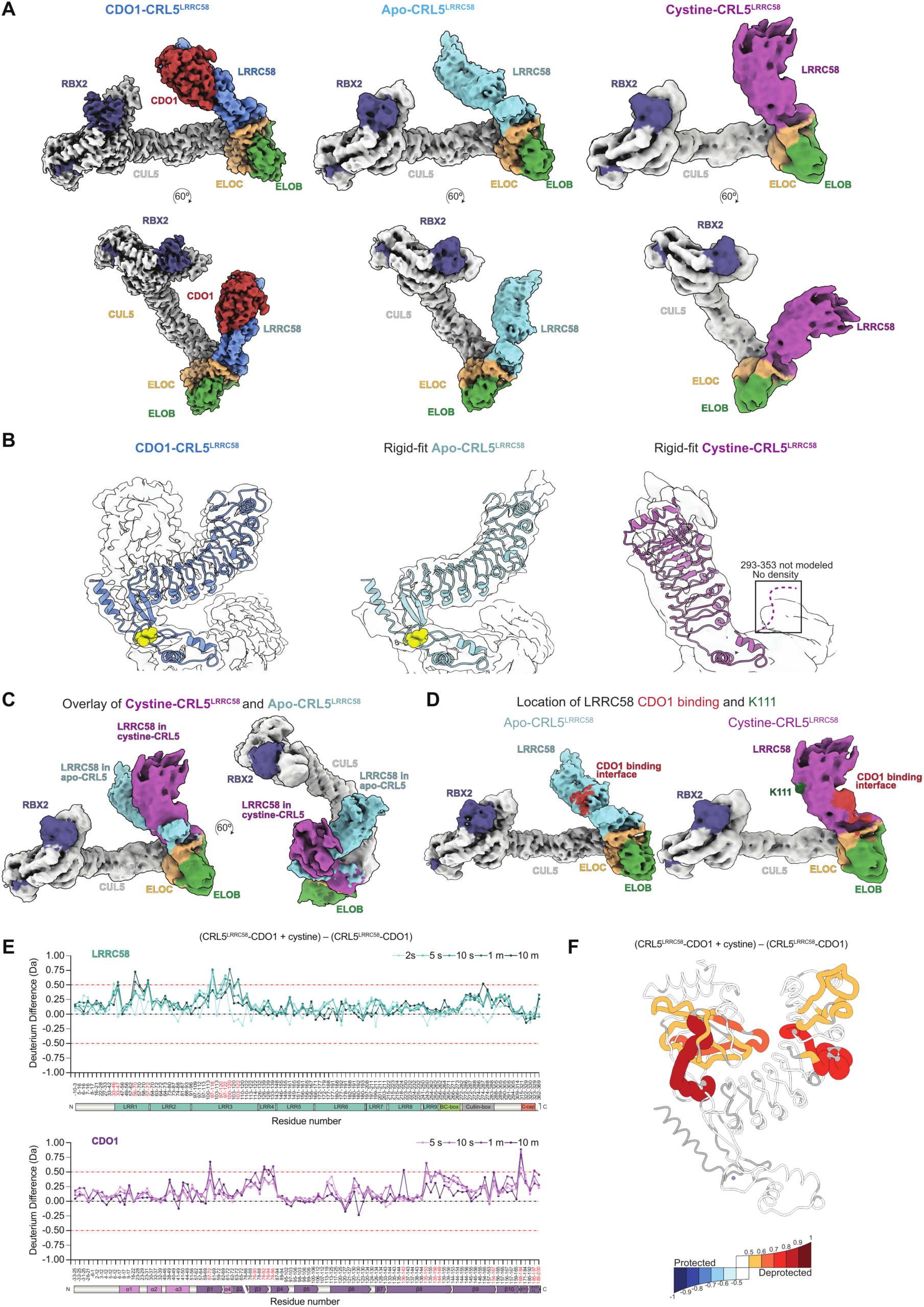
Cyst(e)ine modification of LRRC58 propagates conformational change from the zinc site to the CDO1 interface. **(A)** Comparison of the Cryo-EM maps with LRRC58 colored differently for CDO1-CRL5^LRRC58^(blue), Apo-CRL5^LRRC58^ (light blue), and Cystine-CRL5^LRRC58^ (purple). **(B)** Shown are the fits of the LRRC58 substrate receptor into the corresponding maps of the CDO1-CRL5^LRRC58^ complex, Apo CRL5^LRRC58^, and cystine treated CRL5^LRRC58^. **(C)** Overlay of cystine treated and Apo structure. **(D)** Location of the CDO1 binding region (colored red) and K111 (colored green) in both the Apo and Cystine maps. **(E)** Differential HDX-MS of LRRC58 (top) and CDO1 (bottom) within the CRL5–LRRC58–CDO1 complex, comparing the cystine-treated and untreated states. ΔD = [(CRL5–LRRC58–CDO1 + cystine) − (CRL5–LRRC58–CDO1), Da] versus residue number at the indicated labeling times (2 s–10 min for LRRC58; 5 s–10 min for CDO1; light-to-dark gradient). Data are mean of *n* = 1-4 independent experiments. **(F)** ΔD [(CRL5–LRRC58–CDO1 + cystine) − (CRL5–LRRC58–CDO1)] at the 1 min timepoint mapped onto the cryo-EM structure of the LRRC58-CDO1 subcomplex. Bin edges at ±(0.5, 0.6, 0.7, 0.8, 0.9, and 1.0 Da), saturating at |ΔD| ≥ 1.0 Da. HDX-MS plots: Red dashed lines (±0.5 Da) mark the significance threshold. Domain architecture is annotated below each plot. See “HDX-MS data visualization” section in Materials and Methods.

Moreover, the substrate receptor occupies a different position in the cystine map relative to the CUL5 scaffold and ELOB/ELOC dimer (RMSD of 0.166 Å for the Cullin-ELOB/ELOC), which results in a rotation of the LRR arm of LRRC58 by 38.2 Å and 107.0° relative to the CDO1-bound and Apo structures **(Fig. 3C,D, movie S1)**. The rearrangement is thus confined to LRRC58 and centered on the hinge region connecting the LRR domain with the rest of the complex **(Fig. 3C)**. The 4Cys site staples the base of the loop that anchors the LRR arm to the LRRC58-ELOB/C unit **(Fig. 2C)**. In the CDO1-bound model, this staple is formed by thiolates from both strands of the loop, so that a single zinc clamps the hairpin closed at the point where the arm meets the LRRC58-ELOB/ELOC unit. Covalent modification and zinc ejection at the base are therefore positioned to release a long, substrate-engaging arm. The simplest interpretation of these maps is therefore a directional one: cyst(e)ine drives covalent modification of Cys296 and ejection of structural zinc, the unstapled C-terminal motif loses its fold, and the de-anchored LRR arm rotates as a rigid body. This rotation displaces the substrate receptor by 107.0° corresponding to the rotation of LRRC58, and the spatial relationship between the CDO1 binding interface and RBX2 no longer presents CDO1 productively for ubiquitylation **(Fig. 3D)**.

Consistent with the rigid body rearrangement observed by cryo-EM, HDX-MS of the CRL5^LRRC58^-CDO1 complex revealed cystine-driven deprotection across multiple regions of LRRC58, including the LRR convex face, further demonstrating the conformational change propagates well beyond the Cys296 region of modification **(Fig. 3E,F)**. Similar effects were observed when cystine was added to CRL5^LRRC58^ alone in the absence of CDO1, demonstrating that the restructuring is intrinsic to LRRC58 **(fig. S7A,B)**. In addition, for the CRL5^LRRC58^-CDO1 complex we observed significant cystine-driven deprotection at the CDO1 contact interface **(Fig. 3E,F)**. On CDO1, cystine produced reciprocal deprotection of peptides spanning the LRRC58-binding surface and reversed the CDO1-induced protection at the LRR interface **(Fig. 3E,F)**. These findings illustrate a cystine-driven shift in the dynamic equilibrium between bound and dissociated states, in agreement with the concentration-dependent biochemical disruption by cystine **(Fig. 1)**.

### Cyst(e)ine-driven restructuring of LRRC58 is transmitted via Cys319

Cryo-EM and HDX-MS showed that cyst(e)ine modification of LRRC58 drove conformational changes that include the region of zinc ejection and propagate to rotate the LRR scaffold and reposition its convex and concave faces, while disrupting the CDO1 interaction. Cys319 is positioned along this path, in a loop flanked by the zinc-coordinating cysteine residues including Cys296 and adjacent to the CDO1 binding site **(Fig. 2C)**. This was noteworthy as we found that mutation of Cys319 to alanine or serine abrogates the ability of cysteine to inhibit LRRC58 degradation of CDO1 **(Fig. 2H)**.

To better understand how Cys319 transmits the inhibitory effects driven by cysteine modification of LRRC58, we performed HDX-MS on the CRL5^LRRC58(C319A)^ complex. LRRC58(C319A) retains the capacity to bind CDO1 **(fig. S7C)** and the CRL5^LRRC58(C319A)^ E3 ligase remains constitutively active toward CDO1 **(Fig. 2I)**. HDX-MS revealed that C319A altered baseline LRRC58 dynamics of the LRR convex-facing mobile loop without substantially perturbing the CDO1 binding interface **(fig. S7D-G)**. We next asked how C319A affected the conformational response to cystine. Remarkably, while cystine deprotected the WT LRRC58 CDO1-binding interface and convex LRR mobile loop, these regions did not show significant change in response to cystine in the C319A complex **(Fig. 4A,B).** Therefore, Cys319 is required for the LRR rigid body conformational response of LRRC58 to cystine. Additionally, cystine drives concordant deprotection on the CDO1 side of the interface when incubated with WT LRRC58 **(Fig. 3E,F)**, but with LRRC58(C319A), cystine produces essentially no change to CDO1 **(Fig. 4A,B)**. Moreover, the interface residues between CDO1 and LRRC58 showed minimal differences between the C319A and WT complexes at baseline **(fig. S7D,F)**. Thus, C319A does not lead to tightening of the CDO1 interaction at baseline and specifically diminishes cystine-driven disruption.

**Figure 4.**
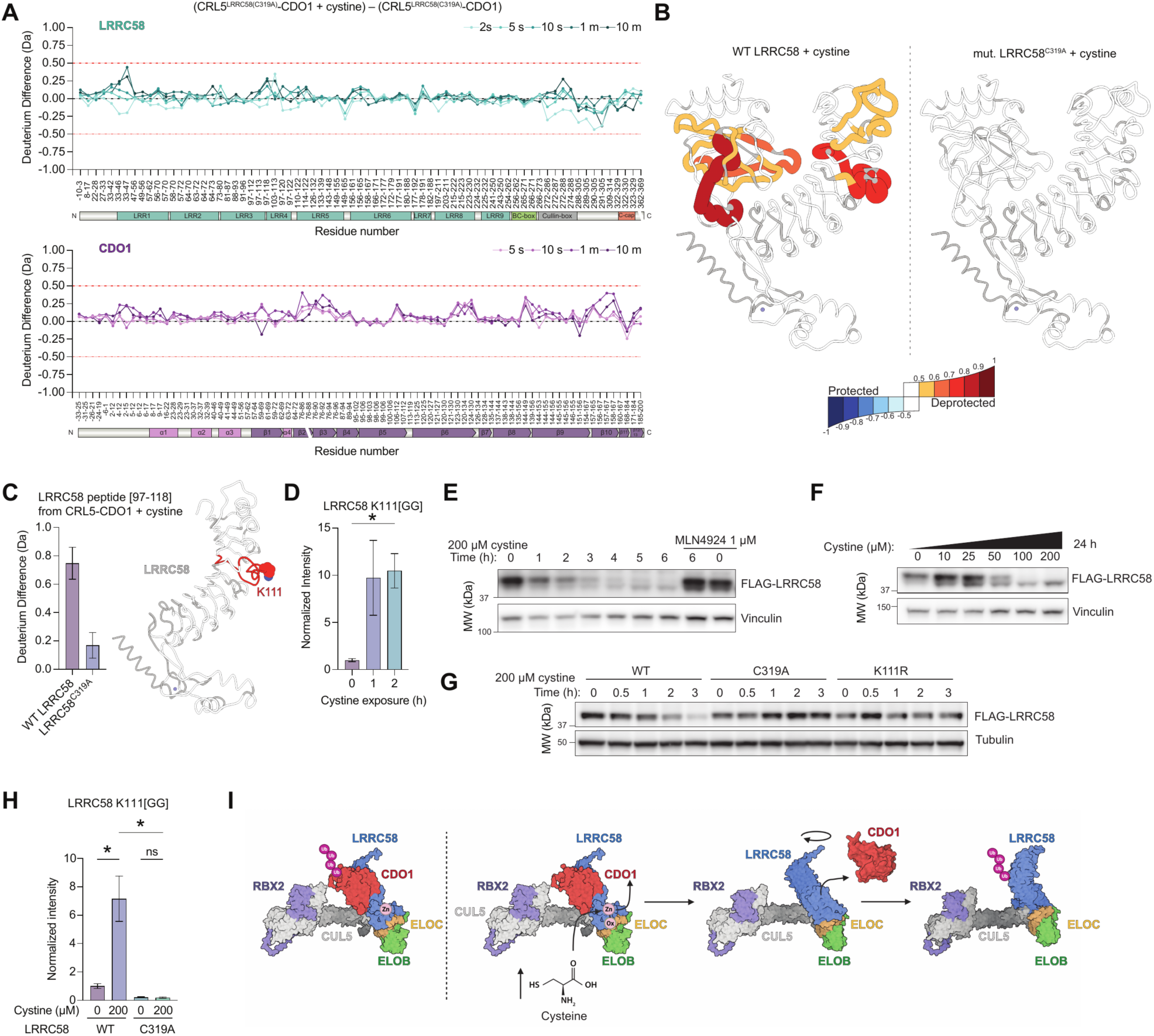
Cys319 relays cyst(e)ine modification to LRRC58 Lys111 to trigger ubiquitylation and degradation. **(A)** Differential HDX-MS of LRRC58 (top) and CDO1 (bottom) within CRL5–LRRC58^C319A^–CDO1, comparing the cystine-treated and untreated states. ΔD = [(mut. CRL5–LRRC58^C319A^–CDO1 + cystine) − (mut. CRL5–LRRC58^C319A^–CDO1), Da] versus residue number at the indicated labeling times (2 s–10 min for LRRC58; 5 s–10 min for CDO1; light-to-dark gradient). Mean of *n* = 2 independent experiments. **(B)** (Left) ΔD [(WT CRL5–LRRC58–CDO1 + cystine) − (WT CRL5–LRRC58–CDO1)] and (Right) ΔD [(mut. CRL5– LRRC58^C319A^–CDO1 + cystine) − (mut. CRL5–LRRC58^C319A^–CDO1)] at the 1 min timepoint mapped onto the cryo-EM structure of the LRRC58-CDO1 subcomplex. Bin edges at ±(0.5, 0.6, 0.7, 0.8, 0.9, and 1.0 Da), saturating at |ΔD| ≥ 1.0 Da. **(C)** Left, Deuterium uptake difference for the LRRC58 97–118 peptide, comparing the cystine-induced change in the WT versus the LRRC58^C319A^ mutant complex. Bars show ΔD for [(WT–CDO1 + cystine) − (WT–CDO1)] and [(C319A–CDO1 + cystine) − (C319A–CDO1)] at the 1 min timepoint, illustrating the loss of cystine-induced deprotection at this peptide in the mutant. *n* = 2 independent experiments. Right, LRRC58 structure with the 97–118 peptide and Lys111 highlighted with corresponding binned color from **(B)**. **(D)** K111[GG] peptide abundance, normalized to two non-cysteine-containing LRRC58 peptides and to the 0 hr condition in cells treated with MG132 plus 200 µM cystine for the indicated duration following overnight cystine depletion. *n* = 5 per group, representative of >4 independent experiments. **(E-F)** Immunoblots of FLAG and vinculin (loading control) in Hep G2 cells overexpressing FLAG-LRRC58 that were cystine-depleted overnight, then treated with cystine for the indicated doses and times, ± 1 μM MLN4924. Each is representative of *n* = 2 independent experiments. **(G)** Immunoblot of FLAG and tubulin (loading control) in Hep G2 LRRC58 KO cells overexpressing FLAG-LRRC58, FLAG-LRRC58 (C319A), or FLAG-LRRC58 (K111R) that were cystine-depleted overnight then treated with 200 μM cystine for the indicated time. Data are representative of *n* = 2 independent experiments. **(H)** Peak area of the K111[GG] peptide, normalized to two non-cysteine-containing LRRC58 peptides, in WT or C319A LRRC58-expressing Hep G2 cells following overnight cystine depletion, MG132 pretreatment, and 4 h MG132 ± 200 µM cystine. *n* = 4–5 per group, representative of 2 independent experiments. **(I)** Cartoon of overall mechanism of cysteine sensing by LRRC58, as supported by the data. Statistics: Data are mean ± SEM unless otherwise noted. Welch’s ANOVA (**D**) and (**H**) with *P*=0.0074 and *P*=0.0034, respectively. Dunnett’s T3 multiple comparisons: * *P*<0.05 as shown. HDX-MS plots: Red dashed lines (±0.5 Da) mark the significance threshold. Domain architecture is annotated below each plot. See “HDX-MS data visualization” section in Materials and Methods.

Consistent with these findings, cystine exhibited a diminished capacity to reduce TR-FRET between LRRC58(C319A)_eGFP_ and CDO1_Tb_ compared to WT proteins, although a residual decrease remained relative to the untreated condition **(fig. S7H)**. The residual TR-FRET response likely reflects the distinct properties measured by the two approaches: HDX-MS resolves solvent protection at specific interface peptides, whereas TR-FRET reports ensemble-averaged fluorophore proximity and may detect rearrangements that do not produce complete interface opening or CDO1 dissociation. Overall, these data are consistent with the cellular phenotype in which CRL5^LRRC58(C319A)^ remains constitutively active towards CDO1 regardless of intracellular cysteine abundance.

Taken together, we show that Cys319 relays modification of the zinc-coordinating cysteine cluster to the LRR domain of LRRC58. Without Cys319, covalent modification does not productively propagate to the CDO1 contact interface and the convex face of LRRC58. LRRC58(C319A) retains CDO1 binding and basal ligase activity but is refractory to cyst(e)ine regulation. The Cys296-containing motif therefore operates as the molecular sensor of cysteine/cystine abundance, and Cys319 transmits this chemical signal into the productive conformational change at the CDO1 interface and the convex face of LRRC58.

### Allosteric action on Cys319 regulates LRRC58 autoubiquitylation

CRL E3 ligase substrate receptors frequently undergo autoubiquitylation, a process by which the E3 ligase conjugates ubiquitin onto its own subunits in addition to, or instead of, recruited substrates^31–36^. These events are thought to be auto-inhibitory and dependent on the accessibility of the target lysine on the substrate receptor to the ubiquitin-charged E2 conjugating enzyme. Based on cryo-EM and HDX-MS, cyst(e)ine modification drives LRR rigid body rotation and presentation of the convex face of LRRC58 toward RBX2 **(Fig. 3D, movie S1)**, positioning a deprotected flexible loop to be a plausible target for ubiquitylation **(Fig. 3E,F)**. Notably, a highly conserved Lys111 centers this flexible loop **(Fig. 4C)**. On this basis, we hypothesized that cystine-driven remodeling of the convex face of the LRRC58 LRR domain could reorient Lys111 to promote ubiquitin conjugation.

To test this, we performed ubiquitylation site mapping by MS on LRRC58 immunoprecipitated from cells subjected to low or high media cystine conditions in order to modify intracellular cysteine concentration^29^. Strikingly, increasing intracellular cyst(e)ine in this way led to a marked increase in di-glycine modification at LRRC58 Lys111, indicative of ubiquitylation at this site **(Fig. 4D & S8A,B)**. Notably, no other detected LRRC58 lysines showed increased di-glycine modification in response to elevated intracellular cyst(e)ine **(fig. S8C)**. Coincident with increased Lys111 ubiquitylation, intracellular cyst(e)ine elevation induced a rapid and concentration-dependent decline in LRRC58 protein levels that was blocked by the neddylation inhibitor MLN4924 **(Fig. 4E,F)**, consistent with previous reports^24,25^. The cyst(e)ine-induced degradation of LRRC58 was no longer seen in LRRC58(K111R) **(Fig. 4G)**. To determine whether cyst(e)ine-driven ubiquitylation of Lys111 was dependent on the Cys319 allosteric relay, we performed the same analysis in LRRC58(C319A) cells. Remarkably, the C319A mutation completely abolished the capacity of cyst(e)ine to promote ubiquitylation at Lys111 **(Fig. 4H)** and decrease LRRC58 protein levels **(Fig. 4G & S8D)**. This result mirrors the HDX-MS findings, in which C319A abolished cystine-driven deprotection of the Lys111-containing convex loop.

Taken together, our data define an allosteric relay through which cyst(e)ine modification of LRRC58 inhibits its ubiquitylation of CDO1 and instead promotes autoubiquitylation **(Fig. 4I)**. Elevated cyst(e)ine drives covalent modification of LRRC58 Cys296 and disruption of the zinc-coordination center that staples a metastable loop containing Cys319. Loss of the zinc staple collapses the loop and propagates through Cys319 to drive rigid-body rotation of the LRR domain which simultaneously disrupts the CDO1 interface while reorienting the convex-facing Lys111 towards the E2 machinery. This rearrangement simultaneously promotes CDO1 dissociation, and LRRC58 autoubiquitylation and degradation, providing the mechanistic basis for a self-contained homeostatic circuit.

### Loss of LRRC58 cysteine sensing leads to systemic cysteine accumulation and early death

To examine the physiological consequences of disabling LRRC58 cysteine sensing, we deployed the C319A LRRC58 substitution in human cells and the orthologous substitution in *C. elegans*. Prime-edited LRRC58(C319A) Hep G2 cells transduced with the CDO1-eGFP reporter **(fig. S5A)** were cultured with ^13^C_6_,^15^N_2_-L-cystine or under cystine restriction. As expected, upon media cystine elevation, LRRC58(C319A) cells exhibited significantly lower levels of CDO1-eGFP compared to WT **(Fig. 5A & S9A)** and failed to effectively catabolize cysteine, exhibiting a near doubling of the total intracellular cysteine pool (**Fig. 5B & S9B**). Moreover, CDO1-mediated catabolism of cysteine to cysteine sulfinic acid (CSA), hypotaurine, and taurine was significantly inhibited in LRRC58^C319A^ cells **(Fig. 5C-E & S9C)**. We also observed elevation in GSH and GSSG production from cysteine in LRRC58(C319A) cells **(Fig. 5F-H & S9D,E)**. Cysteine sensing by LRRC58 is thus required for adaptive control of cellular cysteine concentration in human cells.

**Figure 5.**
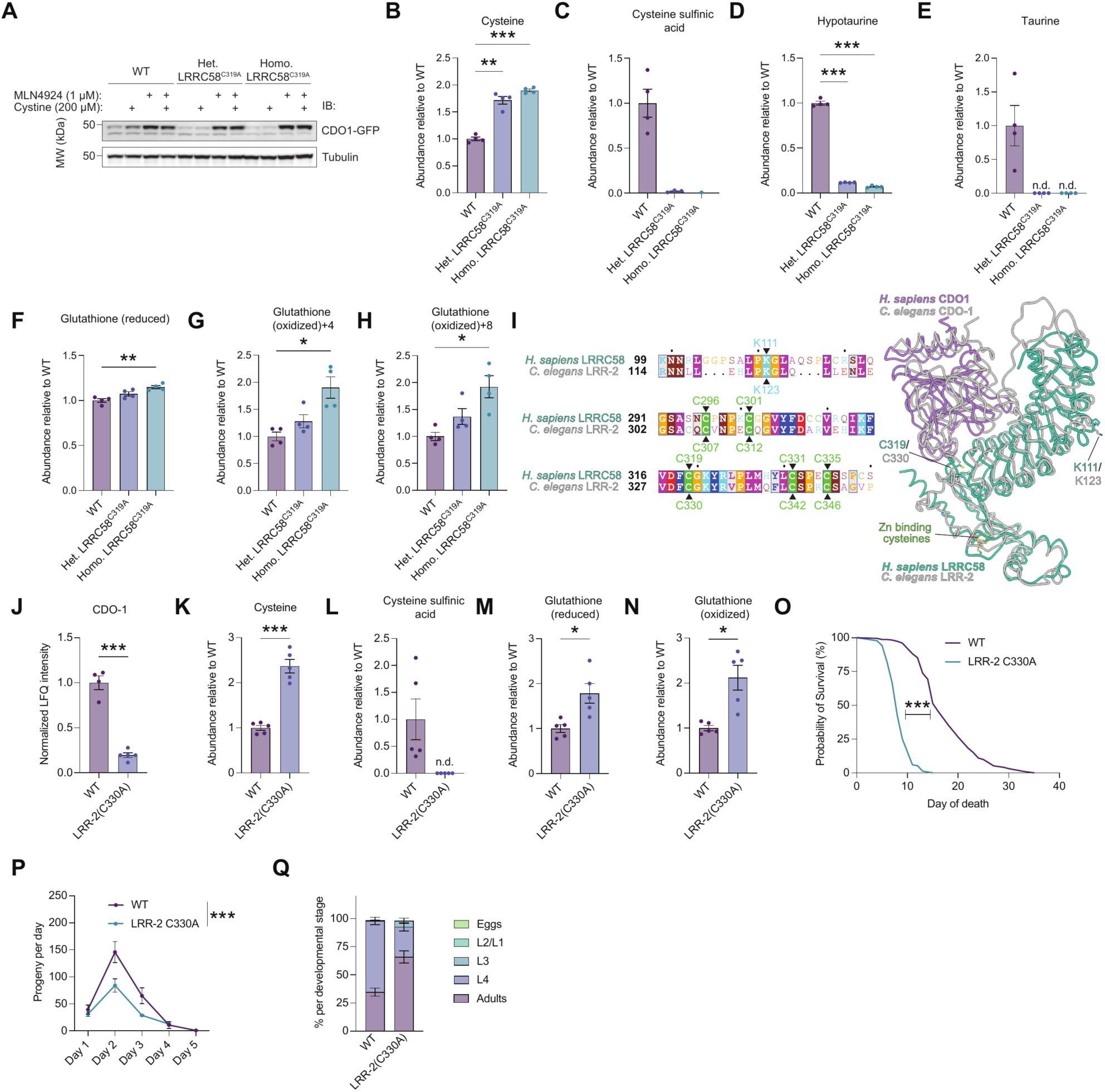
Loss of LRRC58 cysteine-sensing drives cysteine accumulation, organismal toxicity, and early death. **(A)** Immunoblot of CDO1-eGFP and tubulin (loading control) in WT, heterozygous LRRC58^C319A^, and homozygous LRRC58^C319A^ Hep G2 cells, all expressing the CDO1-eGFP reporter (**fig. S5A**). Cells were pretreated with 1 µM MLN4924 or DMSO vehicle for 1 h, then treated for a further 2 h with 200 µM cystine media or cystine-depleted media in the continued presence of 1 µM MLN4924 or DMSO. Data are representative of *n* = 3 biological replicates. **(B-H)** Isotope-tracing metabolomics in WT and CRISPR-engineered LRRC58^C319A^ Hep G2 cells (heterozygous [Het.] and homozygous [Homo.]), expressing the CDO1-eGFP reporter (**fig. S5A**), after 12 h in medium containing ^13^C_6_,^15^N_2_-L-cystine; only tracer-labeled metabolites shown. +4/+8, one/two heavy cysteinyl units incorporated; +3, decarboxylated heavy unit (hypotaurine, taurine). (**B**) Cysteine-NEM+4; (**C**) cysteine sulfinic acid+4; (**D**) hypotaurine+3; (**E**) taurine+3; (**F**) reduced glutathione (GSH)+4; (**G**) oxidized glutathione (GSSG)+4; (**H**) oxidized glutathione (GSSG)+8. *n* = 4 biological replicates. **(I)** (Left) Sequence alignment of the LRR-loop containing Lys111 and the C-terminal region of human LRRC58 and *C. elegans* LRR-2; the ubiquitylated lysine (human Lys111 / worm Lys123), the Zn-binding cysteines (human Cys296 / worm Cys307, human Cys301 / worm Cys312, human Cys331 / worm Cys342, human Cys335 / worm Cys346) and the sensing cysteine (human Cys319 / worm Cys330) are arrowed. (Right) Composite model of the *C. elegans* LRR-2-CDO-1 assembly, generated by superposing AlphaFold3-predicted worm subunits onto the corresponding chains of the human CRL5^LRRC58^-CDO1 cryo-EM structure. The LRR solenoid and C-terminal BC-box domain were superposed independently; dashed line indicates the unmodeled linker. **(J)** CDO-1 protein abundance (normalized LFQ intensity) in WT versus LRR-2(C330A) *C. elegans* by quantitative proteomics (see also **fig. S9F** & **table S2**). *n* = 4 (WT) and *n* = 5 [LRR-2(C330A)] biological replicates. **(K–N)** Steady-state relative abundance of (K) cysteine (NEM-derivatized), (L) CSA, (M) GSH, and (N) GSSG in WT vs. LRR-2(C330A) animals, normalized to WT and to median intensity from paired untargeted metabolomics (worm-number correction). *n* = 5 biological replicates. **(O)** Survival of WT (N2; median 16 days, *n* = 201) and LRR-2(C330A) (median 8 days, *n* = 167) animals at 20 °C. *n* = 2 independent experiments. **(P)** Daily progeny per animal across days 1–5 of adulthood for WT (*n* = 9) and LRR-2(C330A) (*n* = 8) from 3 independent experiments. **(Q)** Developmental stage distribution of synchronized L1 animals scored after 48 h (∼100 L1 per condition), WT versus LRR-2(C330A). *n* = 3 independent experiments. LRR-2(C330A) differed significantly from WT in the Adults (*, *P* = 0.024) and L4 (**, *P* = 0.0057) stages. Statistics: Data are mean ± SEM. Asterisks denote * *P* < 0.05, ** *P* < 0.01, *** *P* < 0.001. Brown-Forsythe and Welch ANOVA with Dunnett’s T3 correction vs. WT, where quantifiable for panels (**B-H**). Unpaired t-test with Welch’s correction vs. WT, where quantifiable for panels (**J**-**N**). Panel level comparisons for panel (**J**) performed on normalized raw intensities. Logrank (Mantel-Cox) test for panel (**O**) and *P* < 0.0001. Mixed-effects analysis (REML) with two-way ANOVA (genotype, day) for panel (**P**), with asterisks denoting a significant main effect of genotype. Unpaired t-tests with Welch’s correction and Holm-Šídák correction for multiple comparisons for panel (**Q**), not depicted on graph for clarity.

*C. elegans* LRR-2 is the ortholog of human LRRC58 and has been shown to regulate worm CDO-1^37^. We found the structural features that mediate the LRRC58-CDO1 interaction, and regulation by cysteine, to be highly conserved between *C. elegans* and human **(Fig. 5I)**. To test the consequences of loss of cysteine sensing *in vivo*, we generated *C. elegans* with an LRR-2 Cys330Ala substitution, corresponding to the cysteine-sensing-deficient human LRRC58(C319A). Quantitative proteomics confirmed CDO-1 protein was constitutively depleted in LRR-2(C330A) animals relative to WT **(Fig. 5J & S9F)**. Moreover, intracellular cysteine more than doubled **(Fig. 5K)**, while the catabolic product of cysteine oxidation by CDO-1 (CSA) was undetectable **(Fig. 5L)**. In parallel, the CDO-1-independent metabolic products of cysteine were elevated in LRR-2(C330A) animals **(Fig. 5M,N)**. Quantitative proteomics of day 1 adult LRR-2(C330A) animals revealed a coherent signature of cysteine overload. Consistent with loss of CDO-1, animals induced RIPS-1, the SAM-dependent methyltransferase that serves as a canonical marker of thiol reductive stress in C. *elegans*^38,39^ **(fig. S9F)** alongside the sulfide-clearance enzyme SQRD-1 and the sulfur-assimilation enzyme CYSL-2 **(fig. S9F)**. This trio constitutes the core HIF-1–dependent response to H_2_S and persulfide accumulation^40–42^, the fate to which excess cysteine is diverted when CDO-1-mediated oxidation is unavailable. LRR-2(C330A) animals also exhibited coordinate loss of glutathione-dependent thiol handling: the SKN-1 target GST-4 was significantly depleted, as was the transsulfuration enzyme CTH-2, consistent with feedback suppression of cysteine synthesis under sustained cysteine excess **(fig. S9F)**.

Because elevated intracellular cysteine drives reductive stress^38^, sulfide accumulation^40^, and promotes iron-dependent Fenton chemistry and cytotoxicity^6,8^, the above proteomic signature reflects a plausible route from failed cysteine sensing to organismal toxicity. Indeed, loss of LRR-2 cysteine sensing in *C. elegans* led to a profound reduction in lifespan. Median survival of LRR-2(C330A) animals was halved to 8 days, compared with 16 days for WT (N2) **(Fig. 5O)**. Additionally, LRR-2(C330A) animals produced nearly half as many progeny as WT **(Fig. 5P)** and had altered developmental progression **(Fig. 5Q)**. Taken together, loss of cysteine sensing uncouples the LRRC58 complex from the cysteine signal, prevents adaptive stabilization of CDO1, and drives systemic cysteine accumulation to manifest reduced fecundity, altered development, and premature death in animals.

## DISCUSSION

Here we describe a molecular mechanism through which cells sense cysteine to maintain intracellular concentrations compatible with healthy life. The CRL5^LRRC58^ E3 ubiquitin ligase uses a cysteine-rich region within its substrate receptor LRRC58 to detect cysteine accumulation through reversible covalent modification and ejection of zinc that staples a metastable loop. This modification propagates through Cys319 to rearrange the substrate-binding interface and expose LRRC58 Lys111 for autoubiquitylation, switching the ligase between CDO1-ubiquitylating and autoubiquitylating states. Cells and animals deprived of this sensor fail to buffer intracellular cysteine and exhibit cysteine accumulation and premature death.

The logic of the LRRC58 mechanism is distinct. Most metabolite homeostasis is indirect: a sensor detects the metabolite, a transcriptional program or signaling cascade relays the signal, and the enzymes of the pathway are adjusted downstream. LRRC58 collapses this architecture into a single complex. Cysteine modifies the receptor directly, and that modification sets the abundance of CDO1, the enzyme that clears it. As such, the metabolite is at once the signal, the modifier of the sensor, and the substrate of the pathway it controls. Because the output is enzyme abundance rather than transcription, the circuit acts post-translationally and the same switch that spares CDO1 commits LRRC58 to its own degradation. That a closed, post-translational circuit has evolved to govern a single metabolite argues that cysteine must be held within narrow limits, and on a timescale that slower forms of regulation cannot meet.

Importantly, cysteine responsiveness can be separated from basal ligase function. Loss of cysteine sensing in human cells and *C. elegans* recapitulates hallmark features of cysteine toxicity. The convergence of these phenotypes downstream of a single substitution at LRRC58 Cys319 establishes that cysteine sensing through LRRC58, not simply the ligase itself, is the rate-limiting variable for protection against cysteine toxicity.

LRRC58 extends an emerging class of redox-and metal-responsive E3 ligases but differs in both the chemistry of the input and the wiring of the response. For example, CUL2^FEM1B^ uses zinc as an intermolecular glue to recognize a reduced FNIP1 degron^43^, whereas LRRC58 coordinates zinc within the receptor itself to maintain an active conformation. KEAP1 likewise uses reactive cysteines to regulate E3 activity but responds to electrophilic and oxidative cues rather than being tuned to cyst(e)ine abundance^44,45^. SCF^FBXL5^ is architecturally similar in coupling metal-dependent structure to substrate recognition and receptor stability^46^, but partitions these functions between distinct iron-containing centers. In LRRC58, a single cysteine-coordinated zinc module couples metabolite sensing to both substrate engagement and receptor fate.

More broadly, these findings suggest that CRL substrate receptors can function as cellular sensors controlling protein degradation. Reactive cysteines, structural metal-binding sites, and related chemical features may provide latent sensing elements that couple metabolites or other cellular cues to substrate recognition and ligase stability. LRRC58 therefore reveals how the ubiquitin system can sense metabolism from within, converting metabolite-responsive chemistry embedded in an E3 ligase directly into adaptive changes in protein stability that maintain homeostasis.

## ACKNOWLEDGEMENTS

The authors thank the HMS Cell Biology Initiative for Genome Editing and Neurodegeneration for the generation of the LRRC58 KO cell line. The authors thank Rui Yan at the HHMI Janelia CryoEM Facility for help in microscope operation and data collection.

## AUTHOR CONTRIBUTIONS

Conceptualization: ECF, ESF, ETC.

Investigation: ECF, HBW, NK, BZ, MAS, YW, JZ, BS, MB, TSL.

Visualization: HBW, NK, ECF. Supervision: TEW, FC, ESF, ETC.

Funding acquisition: ETC, ESF, ECF.

Writing – original draft: ECF, ETC.

Writing – review and editing: all authors.

## DATA AND CODE AVAILABILITY

Cryo-EM density maps have been deposited in the EMDB (accession codes [EMD-79119, EMD-79114, EMD-79115, EMD-79116] for the CDO1-bound, [EMD-79124] for the CDO1-free, and [EMD-79125] for the cystine-modified maps) and the corresponding atomic coordinates in the PDB (PDB: 38SU).

All HDX-MS data referenced are available at massive.ucsd.edu under the data set identifier MSV000102923.

The *C. elegans* proteomics data have been deposited to the ProteomeXchange Consortium via the PRIDE^47^ partner repository with the dataset identifier PXD083730.

The Hep G2 DDA proteomics data for [GG] modifications have been deposited to the ProteomeXchange Consortium via the PRIDE^47^ partner repository with the dataset identifier PXD084432.

Modified Hep G2 cell lines and *C. elegans* strains are available from the corresponding author on request. All remaining data are in the main text or supplementary materials.

## SUPPLEMENTARY MATERIAL

Materials and Methods

Figs. S1 to S9

Tables S1 to S4

Movie S1

Data S1

---

**Figure S1.**
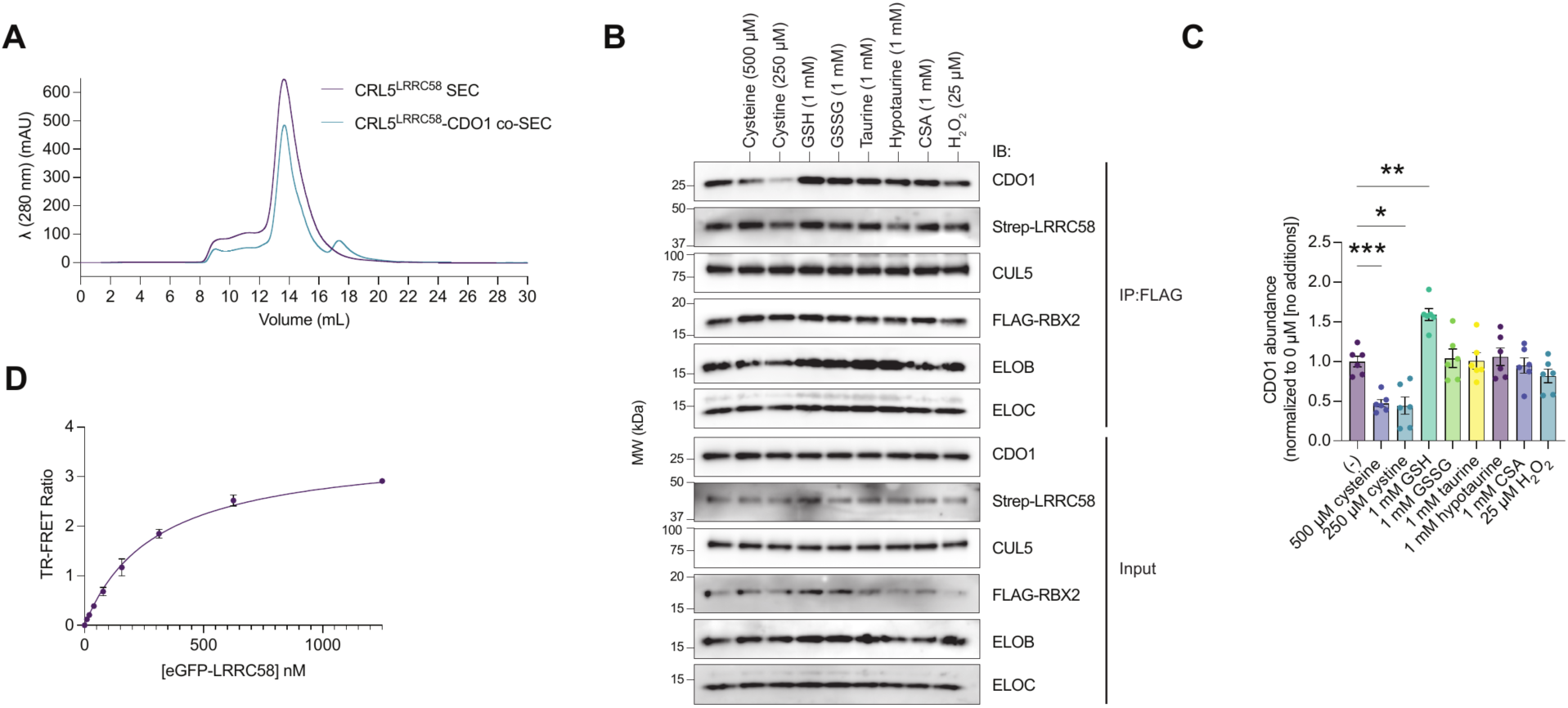
Validation of recombinant CRL5^LRRC58^ and effect of other metabolites on LRRC58-CDO1 interaction. **(A)** SEC trace of co-incubation of CDO1 with CRL5^LRRC58^. **(B)** Representative FLAG IP immunoblot analysis of binding between recombinant preassembled CRL5^LRRC58^ and recombinant Strep∼CDO1 following 1 h preincubation in buffer with various redox-active metabolites related to cellular cysteine metabolism. **(C)** Densitometric quantification of CDO1 abundance from the immunoblots in (**B**). *n* = 5 replicates from 3 independent experiments. **(D)** Time-resolved Förster resonance energy transfer (TR-FRET) signal between LRRC58_eGFP_ and Strep-Avi-CDO1 across increasing concentrations of LRRC58_eGFP_. *n* = 2. Statistics: Data are mean ± SEM. Asterisks denote * *P* < 0.05, ** *P* < 0.01, *** *P* < 0.001. Brown-Forsythe and Welch ANOVA with Dunnett’s T3 correction for panel (**C**).

**Figure S2.**
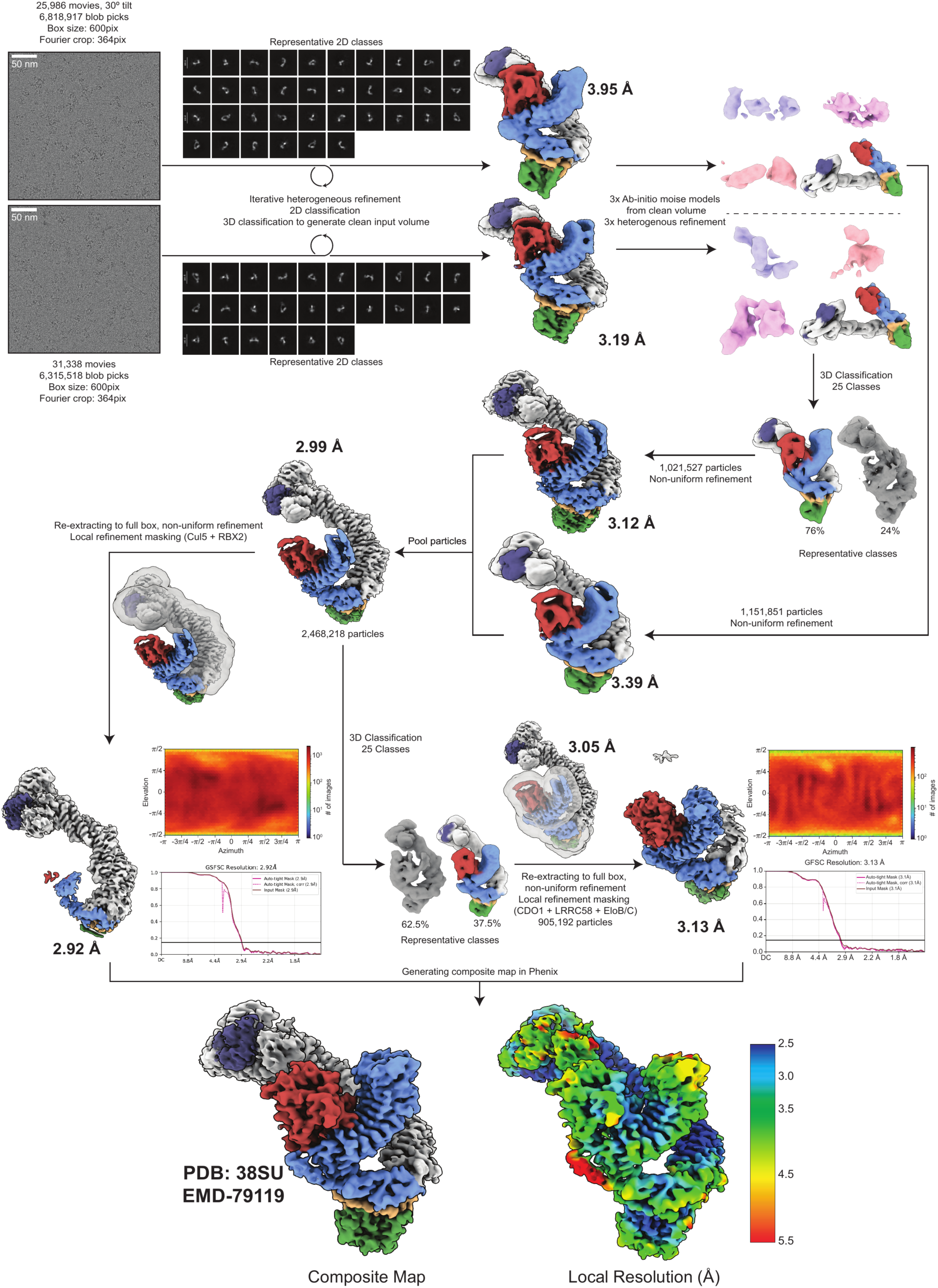
Cryo-EM processing workflow for CRL5^LRRC58^-CDO1 datasets. Overview of processing workflow of two pooled datasets from raw micrograph (low pass filtered to 5 Å) to final composite map of two locally refined maps. The final resolutions of the two locally refined maps shown are post-processing, and the contour level of the final composite map is 12 σ.

**Figure S3.**
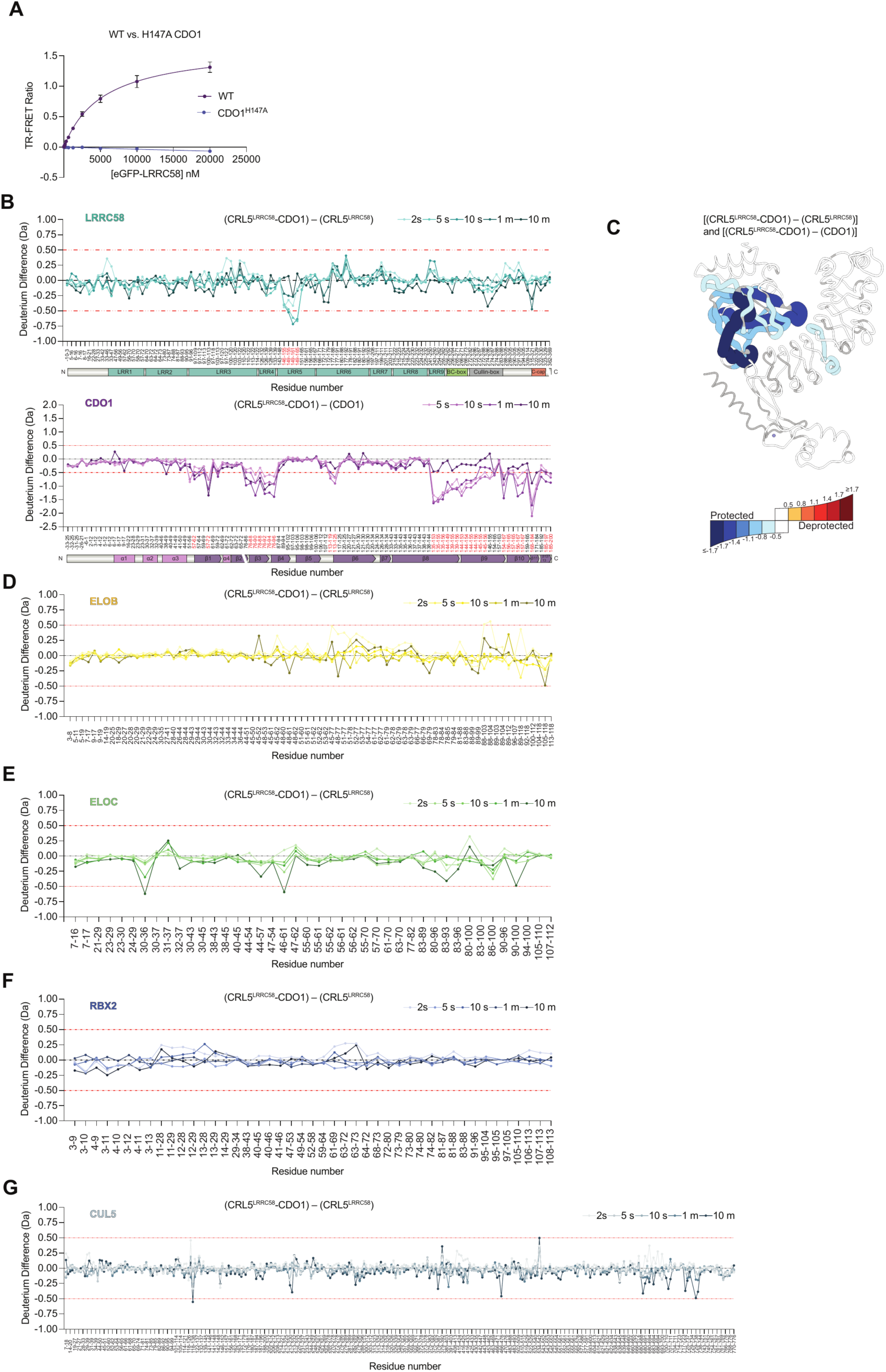
Validation of the LRRC58-CDO1 binding interface by TR-FRET and HDX-MS. **(A)** Time-resolved Förster resonance energy transfer (TR-FRET) signal between LRRC58_eGFP_ and CDO1 or CDO1(H147A) across increasing concentrations of LRRC58_eGFP_. Mean ± SEM of *n* = 2 replicates. **(B)** Differential HDX-MS of LRRC58 (top) and CDO1 (bottom) within the CRL5^LRRC58^–CDO1 complex, comparing the bound and unbound states. ΔD = [(CRL5^LRRC58^–CDO1)-(CRL5^LRRC58^)/(CDO1), Da] versus residue number at the indicated labeling times (2 s–10 min for LRRC58; 5 s–10 min for CDO1; light-to-dark gradient). Data is mean of *n* = 1 or 2 independent experiments. Domain architecture is annotated below each plot. **(C)** ΔD [(CRL5^LRRC58^–CDO1) – (CRL5^LRRC58^)] and ΔD [(CRL5^LRRC58^–CDO1) – (CDO1)] at the 10 s timepoint mapped onto the cryo-EM structure of LRRC58-CDO1 subcomplex. Bin edges at ±(0.5, 0.8, 1.1, 1.4, and 1.7 Da), saturating at |ΔD| ≥ 1.7 Da. **(D-G)** Differential HDX-MS of (**D**) ELOB, (**E**) ELOC, (**F**) RBX2, and (**G**) CUL5 within the CRL5^LRRC58^–CDO1 complex comparing the bound and unbound states. ΔD = [(CRL5^LRRC58^–CDO1) – (CRL5^LRRC58^), Da] versus residue number at the indicated labeling times (2 s–10 min; light-to-dark gradient). Data is mean of *n* = 1 or 2 independent experiments. HDX-MS plots: Red dashed lines (±0.5 Da) mark the significance threshold. See “HDX-MS data visualization” section in Materials and Methods.

**Figure S4.**
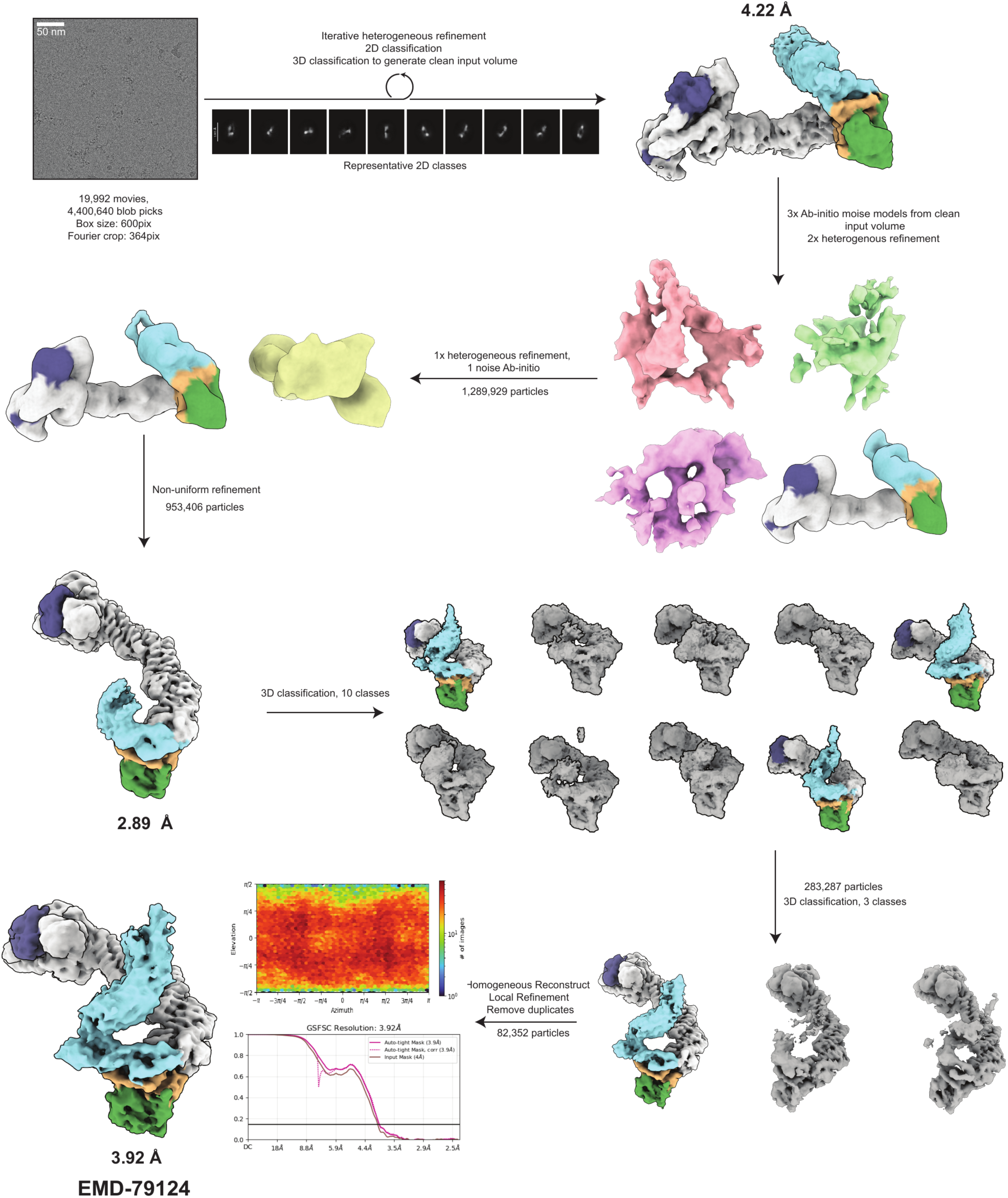
Cryo-EM processing workflow of the Apo-CRL5^LRRC58^ dataset. Overview of processing workflow from raw micrograph (low pass filtered to 5 Å) to final consensus refinement. The final resolution is post-processing, and the contour level of the final map is 12 σ.

**Figure S5.**
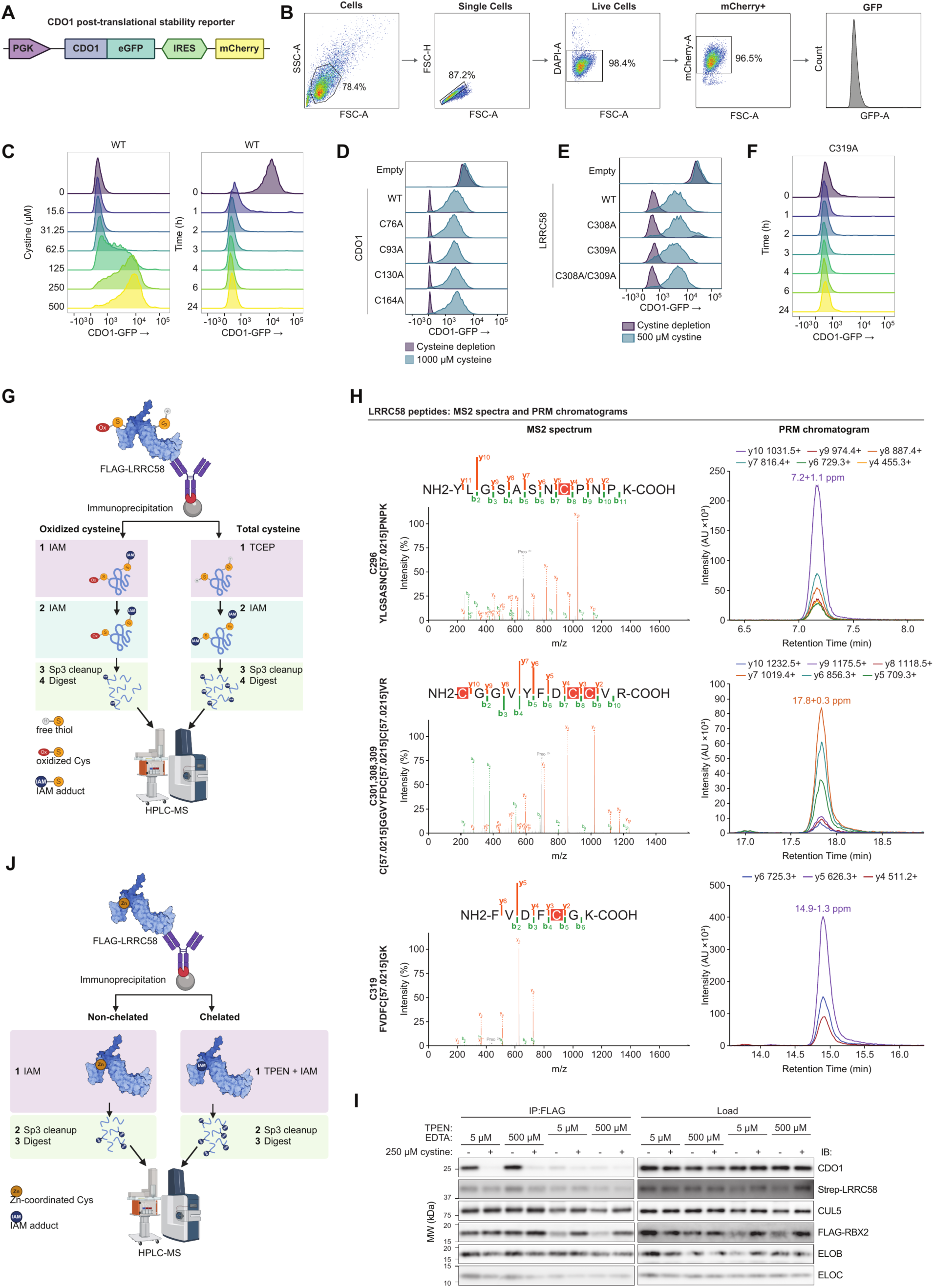
Functional and biochemical characterization of cysteine-dependent regulation of the LRRC58-CDO1 complex. **(A)** Schematic of the lentiviral CDO1 post-translational stability reporter^23^. A phosphoglycerate kinase (PGK) promoter drives expression of a CDO1–enhanced green fluorescent protein (eGFP) C-terminal fusion. An internal ribosome entry site (IRES) allows independent translation of mCherry from the same transcript. **(B)** Flow cytometry gating strategy for cells expressing the CDO1-eGFP reporter. Cells were sequentially gated on forward scatter area (FSC-A) and side scatter area (SSC-A) (cells), FSC-A versus FSC-H (single cells), DAPI exclusion (live cells), and mCherry positivity. CDO1-eGFP fluorescence was quantified within the mCherry⁺ population and are shown as histograms normalized to mode. Percentages indicate the fraction of the parent gate retained at each step for a representative sample in cystine depletion media. **(C-F)** CDO1-eGFP fluorescence (normalized to mode) in the mCherry⁺ population in the indicated cell lines. **(C)** LRRC58 KO Hep G2 cells reconstituted with WT LRRC58 and expressing the WT CDO1 reporter. Left, cells cultured for 24 h in medium containing the indicated concentrations of cystine (0–500 µM). Right, cells cultured in standard media and exposed to cystine depletion media for the indicated times (0–24 h). One representative replicate of 5 independent experiments. **(D)** Hep G2 cells expressing WT or single-cysteine-to-alanine CDO1-eGFP reporters after 24 h in cysteine-depleted (purple) or 1000 µM cysteine (teal) medium. One representative replicate of *n*≥3 technical replicates and 2 independent experiments. **(E)** LRRC58 KO Hep G2 cells expressing the WT CDO1 reporter and reconstituted with empty vector, WT LRRC58, or the indicated single or compound cysteine-to-alanine variants of Cys308 and Cys309, after 24 h in cystine-depleted (purple) or 500 µM cystine (teal) medium. One representative replicate of 2 independent experiments. **(F)** LRRC58 KO Hep G2 cells reconstituted with WT or C319A LRRC58 and expressing the WT CDO1 reporter. Cells were exposed to cystine depletion media for the indicated times (0–24 h). One representative replicate of 2 independent experiments. **(G)** Workflow for quantifying reversible cysteine oxidation on LRRC58 by differential alkylation. FLAG-LRRC58 was immunoprecipitated from Hep G2 cells and each eluate was split into two aliquots. In the “oxidized cysteine” arm, free thiols were capped with iodoacetamide (IAM). In the “total cysteine” arm, all thiols were first reduced with TCEP and then labeled with IAM. Both arms were subjected to SP3 cleanup and on-bead trypsin/Lys-C digestion before HPLC-MS. The fraction of each cysteine that is reversibly oxidized is given by the difference in IAM-labeled peptide signal between the two arms. **(H)** Representative MS2 spectra (left; *m/z* versus relative intensity) and parallel reaction monitoring (PRM) chromatograms (right; retention time versus intensity) for the IAM-labeled (+57.0215 Da) LRRC58 tryptic peptides used for quantification of cysteine oxidation and zinc occupancy, covering C296, C301/C308/C309 (detected as a single peptide), and C319. Annotated b-and y-ion series confirm peptide identity and localize the carbamidomethyl adduct; chromatograms show the transitions detected, with the observed retention time and mass accuracy (ppm) indicated. The most abundant transition was quantified, as shown in **table S4**. **(I)** FLAG IP immunoblot of recombinant preassembled CRL5^LRRC58^ with recombinant Strep-CDO1 after 1 h preincubation with the zinc chelator TPEN (5 to 500 µM) or the comparator chelator EDTA (5 µM to 500 µM), ± cystine. TPEN disrupts the interaction; EDTA has no effect, as expected from their conditional Zn²⁺ affinities and from the inability of EDTA as a tetraanionic hard-donor chelator to access a Zn(Cys)_4_ site. Data are representative of *n* = 2 biological replicates. **(J)** Workflow to measure zinc occupancy at LRRC58 cysteines by differential TPEN chelation. FLAG-LRRC58 was immunoprecipitated from Hep G2 cells. Each eluate was split into two aliquots to be alkylated in the absence (“non-chelated”) or presence (“chelated”) of the zinc chelator TPEN, followed by SP3 cleanup, trypsin/Lys-C digestion, and HPLC-MS. Zinc-coordinated cysteines are protected from alkylation; increased IAM labeling upon TPEN chelation therefore reports the fraction of each cysteine engaged in zinc coordination.

**Figure S6.**
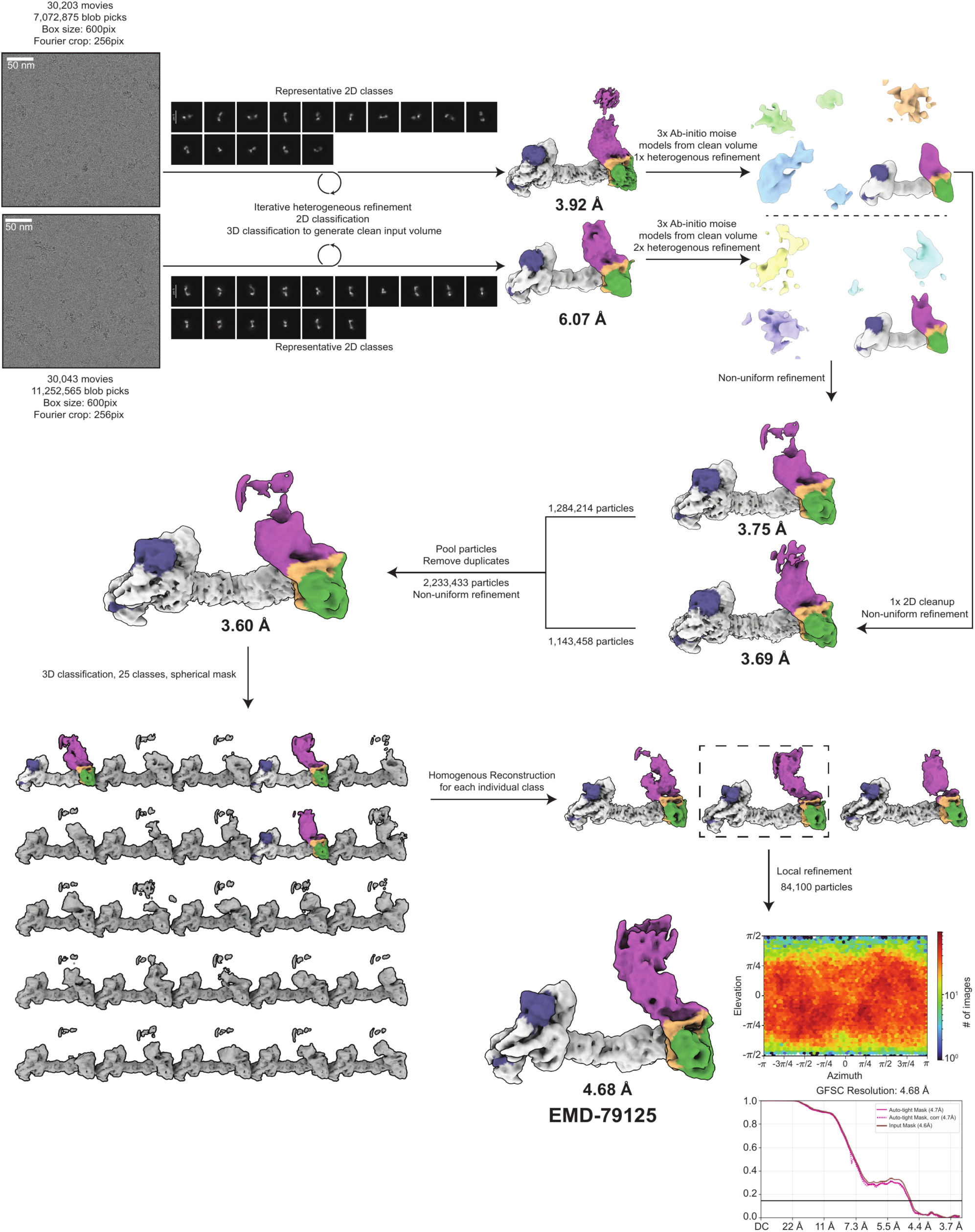
Cryo-EM processing workflow of cystine-CRL5^LRRC58^ datasets. Overview of processing workflow of two pooled datasets from raw micrograph (low pass filtered to 5 Å) to final consensus refinement. The final resolution is post-processing, and the contour level of the final map is 12 σ.

**Figure S7.**
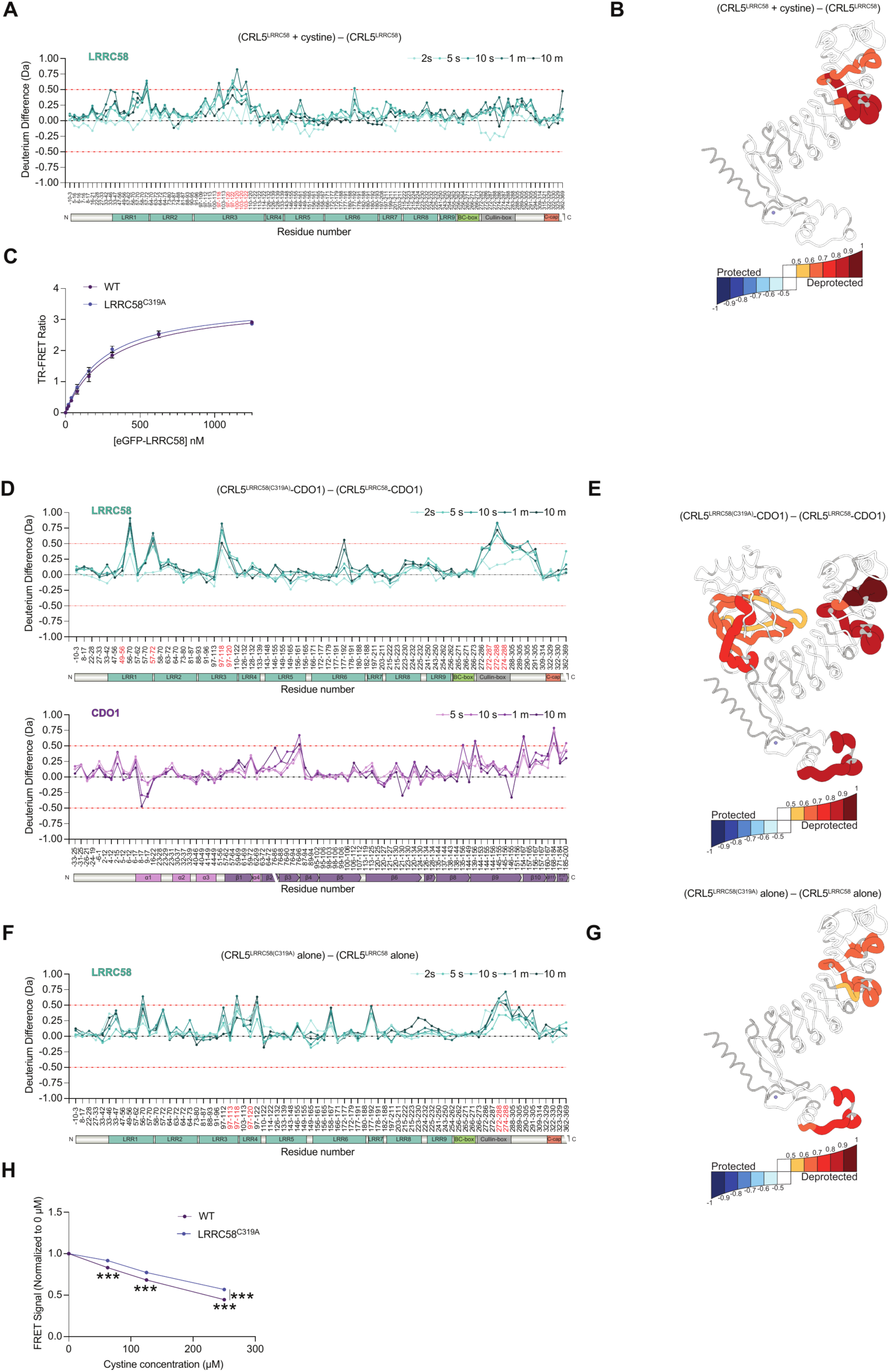
LRRC58(C319A) preserves CDO1 association but alters structural dynamics and cystine responsiveness. **(A)** Differential HDX-MS of LRRC58 within the CRL5^LRRC58^ complex without CDO1, comparing the cystine treated and untreated states. ΔD = [(CRL5–LRRC58 + cystine) - (CRL5–LRRC58), Da] versus residue number at the indicated labeling times (2 s–10 min; light-to-dark gradient). Mean of *n* = 2 independent experiments. **(B)** ΔD [(CRL5–LRRC58 + cystine) - (CRL5–LRRC58)] at the 1 min timepoint mapped onto the cryo-EM structure of Apo-CRL5^LRRC58^ complex (only LRRC58 shown). **(C)** Time-resolved Förster resonance energy transfer (TR-FRET) signal between CDO1 and LRRC58_eGFP_ or LRRC58_eGFP(C319A)_ across increasing concentrations of LRRC58_eGFP_ or LRRC58_eGFP(C319A)_. *n* = 2 replicates; WT data is duplicated from **fig. S1D**. **(D, F)** Differential HDX-MS of LRRC58 (and CDO1, **D** only) within the (**D**) CRL5^LRRC58^–CDO1 complex, or (**F**) the CRL5^LRRC58^ complex alone (lacking CDO1), comparing mutant (CRL5^LRRC58^ ^(C319A)^, ±CDO1) to WT. ΔD = [mutant − WT, Da] vs. residue number at the indicated labeling times [2 s–10 min for LRRC58; 5 s–10 min for CDO1 in (**D**); light-to-dark gradient]. Mean of *n* = 2 independent experiments. **(E, G)** ΔD = [mutant − WT, Da] at the 1 min timepoint mapped onto the cryo-EM structure of (**E**) the LRRC58–CDO1 subcomplex or (**G**) LRRC58 alone, corresponding to the comparisons in (**D**) and (**F**), respectively. **(H)** TR-FRET signal between CDO1 and LRRC58_eGFP_ or LRRC58_eGFP(C319A)_, normalized to 0 µM cystine, across increasing L-cystine concentrations. *n* = 3 replicates, representative of 2 independent experiments. Statistics: Data are mean ± SEM for panels (**C**) and (**H**). *** *P* < 0.001. Two way ANOVA for (**H**), *P* < 0.001 for genotype, cystine dose, and interaction. Sidak’s test for (**H**): P < 0.001, WT vs. LRRC58^C319A^. HDX-MS plots: Red dashed lines (±0.5 Da) mark the significance threshold. Bin edges at ±(0.5–1.0 Da in 0.1 Da increments), saturating at |ΔD| ≥ 1.0 Da. Domain architecture is annotated below each plot. See “HDX-MS data visualization” section in Materials and Methods.

**Figure S8.**
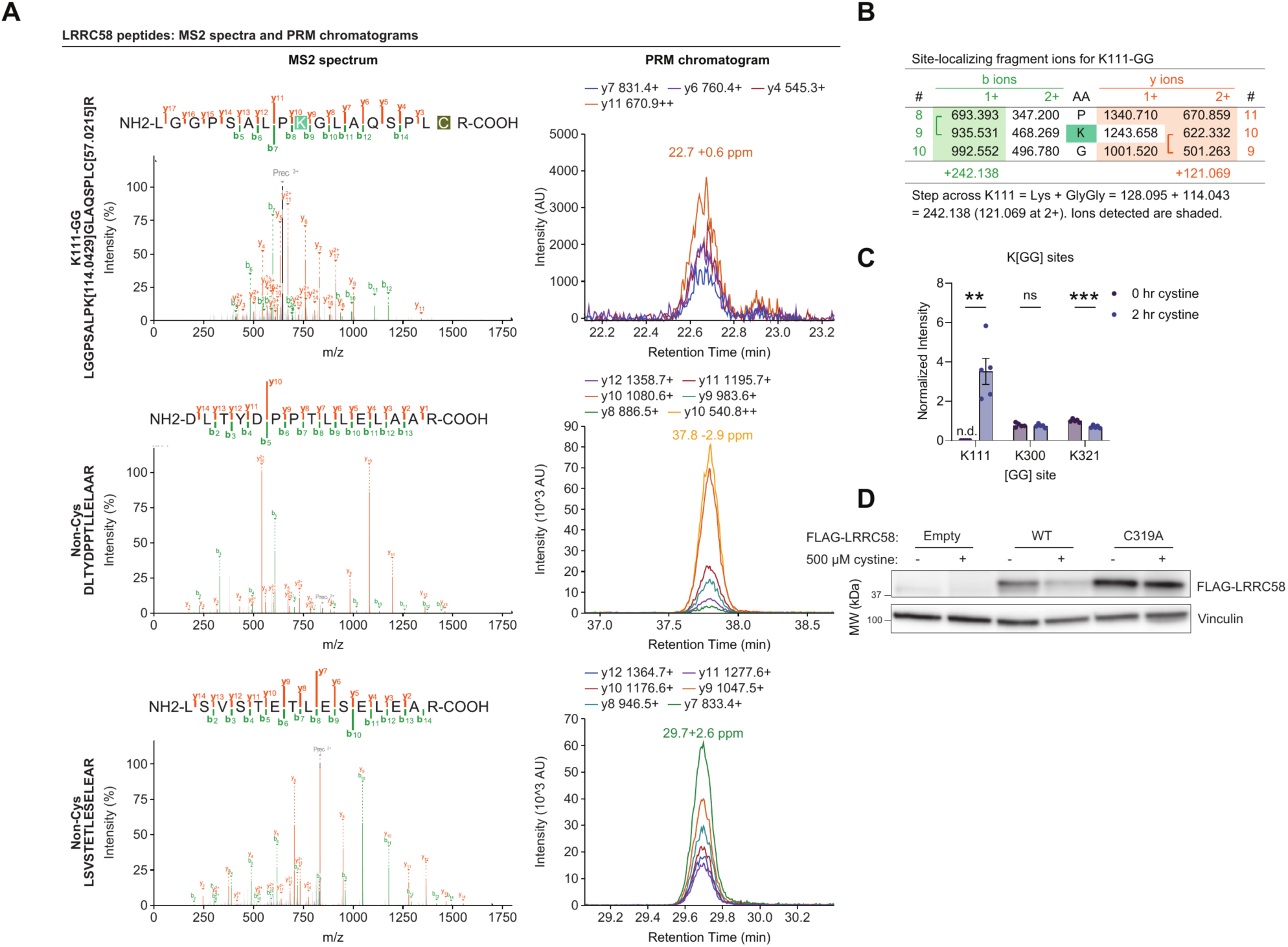
Identification and site localization of diGly-modified LRRC58 Lys111. **(A)** (Left) Representative annotated DDA MS/MS spectrum of the LRRC58 peptide LGGPSALPK[+114.0429]GLAQSPLC[+57.0215]R, containing a diGly[GG] remnant on K111, and two non-cysteine containing LRRC58 peptides used for normalization. For K111[GG], the precursor was detected at *m/z* 646.0187 with charge 3+ and a calibrated precursor mass error of-0.88 ppm. Observed b-type ions are shown in green, y-type ions in orange; superscripts indicate fragment-ion charge states. Surviving precursor is gray. (Right) Extracted PRM fragment-ion chromatograms for various y ions from the LRRC58 K111-diGly and LRRC58 non-cysteine control peptides. The most abundant transition was used for quantitation as listed in table S4. **(B)** Theoretical monoisotopic *m/z* values for fragment ions flanking K111, with detected ions shaded. The b8-to-b9 and y9-to-y10 steps correspond to addition of K[GG], equivalent to 242.138 Da for singly charged ions or 121.069 Th for doubly charged ions. **(C)** Diglycine modification of LRRC58 lysine residues in FLAG-LRRC58 immunoprecipitated from Hep G2 cells following overnight cystine depletion, 1 h MG132 pretreatment, and 2 h continued MG132 ± 200 µM cystine. Mean ± SEM normalized to average intensity; *n* = 5 per group. **(D)** Immunoblot of FLAG or vinculin (loading control) in Hep G2 cells overexpressing empty vector, FLAG-LRRC58 or FLAG-LRRC58 (C319A) that were cystine-depleted overnight and then treated with 500 μM cystine for 24 h. Representative of *n* = 3 independent experiments. Statistics: ** *P* < 0.01, *** *P* < 0.001, ns, not significant. n.d., not detected. Unpaired two-tailed t-tests, corrected by Holm-Šídák for panel (**C**).

**Figure S9.**
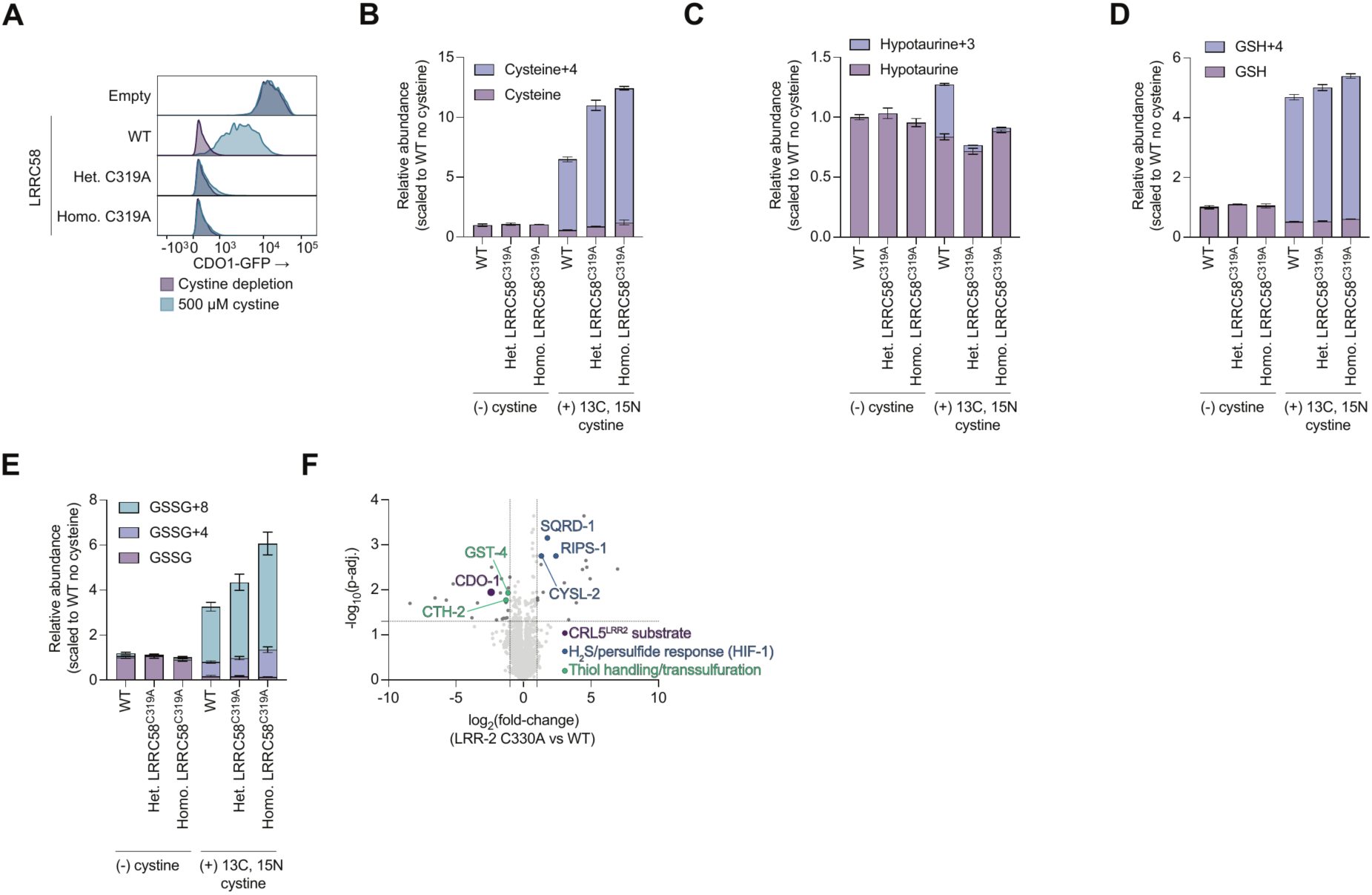
Metabolic and proteomic consequences of disrupting LRRC58 cystine sensing. **(A)** CDO1-eGFP fluorescence normalized to mode in the mCherry^+^ population of WT Hep G2 cells or prime-edited heterozygous (Het.) and homozygous (Homo.) LRRC58^C319A^ Hep G2 cells all expressing the WT CDO1 post-translational stability reporter, after 24 h in cystine-depleted (purple) or 500 µM cystine (teal) medium. Data are representative of *n* = 4 biological replicates. **(B-E)** Total metabolite pools (unlabeled + tracer-derived heavy isotopologues) in WT and CRISPR-engineered LRRC58^C319A^ Hep G2 cells (heterozygous [Het.] and homozygous [Homo.]), all expressing the CDO1-eGFP reporter (**fig. S5A**), measured after 12 h incubation in cystine depleted medium [(-) cysteine] or medium containing 200 µM ^13^C_6_,^15^N_2_-L-cystine [(+) ^13^C, ^15^N cysteine]. Stacked bars show unlabeled (lower) and heavy isotopologue (upper) species; isotopologue superscripts as in Fig. 5 (+4, one heavy cysteinyl unit; +3, decarboxylated unit; +8, two heavy cysteinyl units). Panels show **(B)** cysteine (cysteine/cystine–NEM and cysteine/cystine–NEM+4), (**C**) hypotaurine (hypotaurine and hypotaurine+3), (**D**) reduced glutathione (GSH and GSH+4), and (**E**) oxidized glutathione (GSSG, GSSG+4 and GSSG+8). normalized to WT cystine depletion condition (relative abundance, scaled to WT no cysteine); *n* = 4 biological replicates. **(F)** Volcano plot of quantitative proteomics comparing day-1 adult LRR-2(C330A) and wild-type (N2) *C. elegans*. Log_2_ fold-change (C330A/WT) is plotted against −log_10_ adjusted P-value. Dashed lines mark the significance thresholds (|log_2_ fold-change| > 1, *P*-adj.< 0.05). *n* = 4 [WT] and *n* = 5 [LRR-2(C330A)] biological replicates. Statistics: Data are mean ± SEM for **(B-E)**. *P*-values derived from unpaired t-tests corrected by the Benjamini–Hochberg procedure in panel **(F).**

## REFERENCES

1 Hughes, C. E. et al. Cysteine Toxicity Drives Age-Related Mitochondrial Decline by Altering Iron Homeostasis. Cell 180, 296–310.e218 (2020). 10.1016/j.cell.2019.12.035

2 Nishiuch, Y., Sasaki, M., Nakayasu, M. & Oikawa, A. Cytotoxicity of cysteine in culture media. In Vitro 12, 635–638 (1976). 10.1007/bf02797462

3 Avcilar-Kucukgoze, I. et al. Discharging tRNAs: a tug of war between translation and detoxification in Escherichia coli. Nucleic Acids Res 44, 8324–8334 (2016). 10.1093/nar/gkw697

4 Kumar, A. et al. Homocysteine-and cysteine-mediated growth defect is not associated with induction of oxidative stress response genes in yeast. Biochem J 396, 61–69 (2006). 10.1042/bj20051411

5 Ji, Y. et al. Excessive L-cysteine induces vacuole-like cell death by activating endoplasmic reticulum stress and mitogen-activated protein kinase signaling in intestinal porcine epithelial cells. Amino Acids 48, 149–156 (2016). 10.1007/s00726-015-2071-5

6 Nakamura, T. et al. Cysteine excess triggers a mitochondrial iron-dependent cell death. Nature Metabolism (2026). 10.1038/s42255-026-01616-7

7 Brain, J. A. et al. Excess cysteine drives conjugate formation and impairs proliferation of NRF2-activated cancer cells. Nature Metabolism 8, 840–854 (2026). 10.1038/s42255-026-01499-8

8 Park, S. & Imlay, J. A. High levels of intracellular cysteine promote oxidative DNA damage by driving the fenton reaction. J Bacteriol 185, 1942–1950 (2003). 10.1128/JB.185.6.1942-1950.2003

9 Janáky, R., Varga, V., Hermann, A., Saransaari, P. & Oja, S. S. Mechanisms of L-cysteine neurotoxicity. Neurochem Res 25, 1397–1405 (2000). 10.1023/a:1007616817499

10 Puka-Sundvall, M., Eriksson, P., Nilsson, M., Sandberg, M. & Lehmann, A. Neurotoxicity of cysteine: interaction with glutamate. Brain Research 705, 65–70 (1995). 10.1016/0006-8993(95)01139-0

11 Pedersen, O. & Karlsen, R. L. The toxic effect of L-cysteine on the rat retina. A morphological and biochemical study. Investigative Ophthalmology & Visual Science 19, 886–892 (1980).

12 Shibui, Y., Sakai, R., Manabe, Y. & Masuyama, T. Comparisons of l-cysteine and d-cysteine toxicity in 4-week repeated-dose toxicity studies of rats receiving daily oral administration. J Toxicol Pathol 30, 217–229 (2017). 10.1293/tox.2017-0002

13 Bradley, H. et al. Sulfate metabolism is abnormal in patients with rheumatoid arthritis. Confirmation by in vivo biochemical findings. The Journal of rheumatology 21, 1192–1196 (1994).

14 Heafield, M. T. et al. Plasma cysteine and sulphate levels in patients with motor neurone, Parkinson’s and Alzheimer’s disease. Neuroscience Letters 110, 216–220 (1990). 10.1016/0304-3940(90)90814-P

15 Özkan, Y., Özkan, E. & Şimşek, B. Plasma total homocysteine and cysteine levels as cardiovascular risk factors in coronary heart disease. International Journal of Cardiology 82, 269–277 (2002). 10.1016/S0167-5273(02)00010-4

16 Jacob, N., Bruckert, E., Giral, P., Foglietti, M. J. & Turpin, G. Cysteine is a cardiovascular risk factor in hyperlipidemic patients. Atherosclerosis 146, 53–59 (1999). 10.1016/s0021-9150(99)00128-8

17 Li, J. et al. Multimodal clocks of human aging. Cell 189, 4489–4511.e4431 (2026). 10.1016/j.cell.2026.04.025

18 Abu-Remaileh, M. et al. Lysosomal metabolomics reveals V-ATPase-and mTOR-dependent regulation of amino acid efflux from lysosomes. Science 358, 807–813 (2017). 10.1126/science.aan6298

19 Roman, H. B. et al. The Cysteine Dioxgenase Knockout Mouse: Altered Cysteine Metabolism in Nonhepatic Tissues Leads to Excess H2S/HS− Production and Evidence of Pancreatic and Lung Toxicity. Antioxidants & Redox Signaling 19, 1321–1336 (2013). 10.1089/ars.2012.5010

20 Bella, D. L., Hirschberger, L. L., Hosokawa, Y. & Stipanuk, M. H. Mechanisms involved in the regulation of key enzymes of cysteine metabolism in rat liver in vivo. American Journal of Physiology-Endocrinology and Metabolism 276, E326–E335 (1999).

21 Bella, D. L., Hahn, C. & Stipanuk, M. H. Effects of nonsulfur and sulfur amino acids on the regulation of hepatic enzymes of cysteine metabolism. American Journal of Physiology-Endocrinology and Metabolism 277, E144–E153 (1999).

22 Stipanuk, M. H., Hirschberger, L. L., Londono, M. P., Cresenzi, C. L. & Yu, A. F. The ubiquitin-proteasome system is responsible for cysteine-responsive regulation of cysteine dioxygenase concentration in liver. American Journal of Physiology-Endocrinology and Metabolism 286, E439–E448 (2004).

23 Xiao, H. et al. Covariation MS uncovers a protein that controls cysteine catabolism. Nature (2025). 10.1038/s41586-025-09535-5

24 Ramage, D. E. et al. LRRC58 defines an E3 ubiquitin ligase complex sensitive to cysteine abundance. Molecular Cell 86, 2828–2842.e2812 (2026). 10.1016/j.molcel.2026.06.014

25 Andree, G. A. et al. Cysteine availability tunes ubiquitin signaling via inverse stability of LRRC58 E3 ligase and its substrate CDO1. Nature Communications 17, 4196 (2026). 10.1038/s41467-026-72524-3

26 Sober, E. Ockham’s Razors: A User’s Manual. (Cambridge University Press, 2015).

27 Ziegler, D. M. Role of reversible oxidation-reduction of enzyme thiols-disulfides in metabolic regulation. Annu Rev Biochem 54, 305–329 (1985). 10.1146/annurev.bi.54.070185.001513

28 Hochgrafe, F. et al. S-cysteinylation is a general mechanism for thiol protection of Bacillus subtilis proteins after oxidative stress. J Biol Chem 282, 25981–25985 (2007). 10.1074/jbc.C700105200

29 Conrad, M. & Sato, H. The oxidative stress-inducible cystine/glutamate antiporter, system xc−: cystine supplier and beyond. Amino Acids 42, 231–246 (2012). 10.1007/s00726-011-0867-5

30 Burger, N. et al. The human zinc-binding cysteine proteome. Cell 188, 832–850 e827 (2025). 10.1016/j.cell.2024.11.025

31 Sugasawa, K. et al. UV-induced ubiquitylation of XPC protein mediated by UV-DDB-ubiquitin ligase complex. Cell 121, 387–400 (2005). 10.1016/j.cell.2005.02.035

32 Groisman, R. et al. CSA-dependent degradation of CSB by the ubiquitin-proteasome pathway establishes a link between complementation factors of the Cockayne syndrome. Genes Dev 20, 1429–1434 (2006). 10.1101/gad.378206

33 Zhou, C. et al. Fission yeast COP9/signalosome suppresses cullin activity through recruitment of the deubiquitylating enzyme Ubp12p. Mol Cell 11, 927–938 (2003). 10.1016/s1097-2765(03)00136-9

34 Bornstein, G., Ganoth, D. & Hershko, A. Regulation of neddylation and deneddylation of cullin1 in SCFSkp2 ubiquitin ligase by F-box protein and substrate. Proc Natl Acad Sci U S A 103, 11515–11520 (2006). 10.1073/pnas.0603921103

35 Schmidt, M. W., McQuary, P. R., Wee, S., Hofmann, K. & Wolf, D. A. F-box-directed CRL complex assembly and regulation by the CSN and CAND1. Mol Cell 35, 586–597 (2009). 10.1016/j.molcel.2009.07.024

36 Bennett, E. J., Rush, J., Gygi, S. P. & Harper, J. W. Dynamics of cullin-RING ubiquitin ligase network revealed by systematic quantitative proteomics. Cell 143, 951–965 (2010). 10.1016/j.cell.2010.11.017

37 Patange, O., Breen, P., Arsuffi, G. & Ruvkun, G. Hydrogen sulfide mediates the interaction between C. elegans and Actinobacteria from its natural microbial environment. Cell Rep 44, 115170 (2025). 10.1016/j.celrep.2024.115170

38 Ravi, Kumar, A., Bhattacharyya, S. & Singh, J. Thiol reductive stress activates the hypoxia response pathway. EMBO J 42, e114093 (2023). 10.15252/embj.2023114093

39 Winter, A. D. et al. Dietary-derived vitamin B12 protects Caenorhabditis elegans from thiol-reducing agents. BMC Biol 20, 228 (2022). 10.1186/s12915-022-01415-y

40 Budde, M. W. & Roth, M. B. The response of Caenorhabditis elegans to hydrogen sulfide and hydrogen cyanide. Genetics 189, 521–532 (2011). 10.1534/genetics.111.129841

41 Miller, D. L., Budde, M. W. & Roth, M. B. HIF-1 and SKN-1 coordinate the transcriptional response to hydrogen sulfide in Caenorhabditis elegans. PLoS One 6, e25476 (2011). 10.1371/journal.pone.0025476

42 Ma, D. K., Vozdek, R., Bhatla, N. & Horvitz, H. R. CYSL-1 interacts with the O2-sensing hydroxylase EGL-9 to promote H2S-modulated hypoxia-induced behavioral plasticity in C. elegans. Neuron 73, 925–940 (2012). 10.1016/j.neuron.2011.12.037

43 Manford, A. G. et al. Structural basis and regulation of the reductive stress response. Cell 184, 5375–5390.e5316 (2021). 10.1016/j.cell.2021.09.002

44 Dinkova-Kostova, A. T. et al. Direct evidence that sulfhydryl groups of Keap1 are the sensors regulating induction of phase 2 enzymes that protect against carcinogens and oxidants. Proc Natl Acad Sci U S A 99, 11908–11913 (2002). 10.1073/pnas.172398899

45 Zhang, D. D. & Hannink, M. Distinct cysteine residues in Keap1 are required for Keap1-dependent ubiquitination of Nrf2 and for stabilization of Nrf2 by chemopreventive agents and oxidative stress. Mol Cell Biol 23, 8137–8151 (2003). 10.1128/MCB.23.22.8137-8151.2003

46 Wang, H. et al. FBXL5 Regulates IRP2 Stability in Iron Homeostasis via an Oxygen-Responsive [2Fe2S] Cluster. Mol Cell 78, 31–41.e35 (2020). 10.1016/j.molcel.2020.02.011

47 Perez-Riverol, Y. et al. The PRIDE database at 20 years: 2025 update. Nucleic Acids Res 53, D543–D553 (2025). 10.1093/nar/gkae1011

