## Supplementary Information for "The mechanism of cysteine sensing by LRRC58"

**The PDF file includes:**

Materials and Methods
Figs. S1 to S9
References (*48-60*)

**Other Supplementary Materials for this manuscript include the following:**

Movie S1
Tables S1-S4
Data S1

### Materials and Methods

**Cell lines and culture conditions.** Hep G2 cells were obtained from the American Type Culture Collection (ATCC) and maintained in Eagle's Minimum Essential Medium (EMEM; ATCC, 30-2003) supplemented with 10% fetal bovine serum (FBS; GeminiBio, 100-106) and 1% penicillin-streptomycin (Corning, 30-002-CI). Cells were detached with 0.25% trypsin–EDTA (Gibco, 25200-056) and passaged every 3 to 4 days. Cells were maintained at 37°C in 5% $CO_2$. Cultures were tested for mycoplasma contamination and were negative.

Lenti-X 293T cells (Takara Bio, 632180) used for lentiviral production were maintained in Dulbecco's Modified Eagle Medium (DMEM, Corning 10-017-CV) supplemented with 10% FBS and 1% penicillin–streptomycin.

Sf9 cells were obtained from Gibco and were cultured in ESF 921 medium (Expression Systems). High Five cells were obtained from Gibco and were cultured in SF-4 Baculo Express ICM medium (BioConcept) or ExpiSF-CD medium (Gibco). Both were grown at 27°C with 120rpm shaking.

**Reagents.** MG132 (MedChemExpress, HY-13259 or Selleck Chem S2619), MLN4924 (MedChemExpress, HY-70062), L-cystine (Thermo Scientific, J61651.09), $^{13}C_6$,$^{15}N_2$-L-cystine (Cambridge Isotope Laboratories, CNLM-4244-H-0.1), L-cysteine (Thermo Scientific J63745), L-methionine (Thermo Fisher, J61904.18), L-cystine dihydrochloride (Sigma Aldrich C6727), TPEN (N,N,N',N'-Tetrakis(2-pyridylmethyl)ethylenediamine, Sigma-Aldrich, P4413), TCEP (Bond Breaker, Thermo Scientific, 77720), DTT (Thermo Fisher, A39255), iodoacetamide (I1149, Sigma Aldrich), N-ethylmaleimide (Thermo Fisher, 23030), guanidine hydrochloride (Thermo Fisher, A13543.0B), Benzonase Nuclease (Sigma-Aldrich/Novagen, 70664-3), ammonium bicarbonate (Sigma-Aldrich, 09830), kanamycin sulfate (Sigma-Aldrich, K4000), 5-fluoro-2'-deoxyuridine (Thermo Scientific, L16497.ME), polybrene (Santa Cruz, sc-134220), puromycin dihydrochloride (Thermo Fisher, A1113803), blasticidin S (Thermo Fisher, A1113903), Lipofectamine 3000 (Thermo Fisher, L3000015), TransIT-LT1 (Mirus, MIR2300), DAPI (Cell Signaling Technology, 4083S), anti-FLAG M2 magnetic beads (Sigma-Aldrich, M8823), 3X FLAG peptide (Sigma-Aldrich, F4799), Tb-streptavidin (terbium: FischerSci 69705, streptavidin: Genscript Z02043-5), SP3 carboxylate-modified magnetic beads (Cytiva, 1:1 mix of 45152105050350 and 65152105050350), sequencing-grade modified trypsin (Promega, V5113), Lys-C (Fisher Scientific, NC9309054), protease inhibitor tablets (Roche, 11836153001), BCA protein assay kit (Thermo Fisher, 23225), LC-MS-grade methanol, acetonitrile and formic acid (Fisher Scientific, A456-4, A955-4 and A11750), and Evotip Pure tips (Bruker, Q-98763, 1897435) were used throughout. Antibodies and their dilutions are listed under **Western blotting**. Reagents are referred to by name only in the sections below.

**Lentiviral constructs.** The previously described CDO1 post-translational stability reporter consists of full-length human CDO1 fused to enhanced green fluorescent protein (eGFP), followed by an internal ribosome entry site and mCherry within the Cilantro 2 lentiviral vector[23]. Cilantro 2 was a gift from B. Ebert (Addgene plasmid 74450; RRID: Addgene_74450). CDO1 point mutants were synthesized and cloned into the same vector by Twist Bioscience. Wild-type and mutant FLAG-tagged LRRC58 constructs were synthesized and cloned into pTwist Lenti CMV BSD by Twist Bioscience. The FLAG tag was positioned at the N-terminus of LRRC58. All constructs were verified by whole-plasmid sequencing. The complete list of constructs and mutations is provided in **table S3**.

**Lentivirus production and transduction.** Lentivirus was produced in Lenti-X 293T cells by cotransfection of the lentiviral expression vector with psPAX2 (Addgene #12260; RRID: Addgene_12260) and the VSV-G envelope plasmid pMD2.G (Addgene #12259; RRID: Addgene_12259), both gifts from Didier Trono, using TransIT-LT1. Virus-containing medium was collected 48 h after transfection and passed through a 0.45 μm filter.

Hep G2 cells were transduced by centrifugation-assisted infection in the presence of polybrene. Medium was replaced 24–48 h after transduction. Beginning 24-48 hours after transduction, cells were selected with puromycin or blasticidin.

**Generation of LRRC58-knockout Hep G2 cells.** Hep G2 cells were cultured in DMEM/F-12 medium (Gibco, 11320033) supplemented with 10% FBS (Corning). The HiFi Cas9 expression plasmid was generated by introducing a R691A mutation into the Cas9 coding sequence in pET-Cas9-NLS-6xHis (Addgene plasmid # 62933 ref[48]). HiFi Cas9 protein was expressed in Rosetta™(DE3)pLysS Competent Cells (Novagen, 70956) and purified as described elsewhere[49]. To generate LRRC58 knockout Hep G2 cells, the sgRNA targeting the sequence 5' ACCGTCTGGTGTCGCTGCCA 3' was generated using GeneArt Precision gRNA Synthesis Kit (Thermo Fisher Scientific, A29377). Transfection was performed with the Neon Transfection System (Thermo Fisher Scientific). Briefly, 0.6 μg sgRNA was incubated with 3 μg HiFi Cas9 protein for 10 minutes at room temperature and electroporated into 2 x $10^5$ Hep G2 cells. Transfected cells were seeded at clonal density and colonies were picked after one week. Knockouts were identified by Illumina MiSeq sequencing. Briefly, the library was prepared using a two-step PCR approach. The primers for amplicon PCR were 5'-ACACTCTTTCCCTACACGACGCTCTTCCGATCTCGAGACGCTGGAGTCTGAG and 5'-GTGACTGGAGTTCAGACGTGTGCTCTTCCGATCTGGTTGTTCTTGGCCAGCAG. The primers for index PCR were 5'-AATGATACGGCGACCACCGAGATCTACAC$(n)_{10}$ACACTCTTTCCCTACACGACGCTC and 5'- CAAGCAGAAGACGGCATACGAGAT$(n)_{10}$ GTGACTGGAGTTCAGACGTGTGCT. Aliquots of PCR product from each sample were pooled and purified using a Qiagen PCR purification kit. The library was paired end sequenced on an Illumina MiSeq according to the standard protocols (Illumina).

Two clones were primarily used in the experiments presented here, with sequencing showing indels which were predicted to be damaging (40 nucleotide (nt) deletion/8 nt deletion and 20 nt deletion/11 nt deletion). The functional behavior of these cell lines was further confirmed by lack of effect of cystine on the CDO1 post-translational stability reporter in the absence of LRRC58 complementation.

**Generation of prime-edited LRRC58(C319A) Hep G2 cells.** The C319A substitution was introduced into Hep G2 cells by prime editing using the PE3 strategy, performed essentially as described in ref. [50,51] with the PEmax editor architecture[52]. An epegRNA encoding the C319A edit (spacer, primer-binding site, reverse-transcription template, and 3' tevopreQ1 motif) and a PE3 nicking sgRNA were designed following the guidelines in ref.[50] and cloned by Golden Gate assembly into pU6-tevopreq1-GG-acceptor (Addgene #174038) and pU6-pegRNA-GG-acceptor (Addgene #132777), respectively; spacer, PBS, RTT, and nicking-sgRNA sequences are listed in **table S3**. Hep G2 cells were co-transfected with pCMV-PEmax-P2A-GFP (Addgene #180020), the epegRNA, and the nicking-sgRNA plasmids using Lipofectamine 3000. 72 h after transfection, GFP-positive cells were isolated by FACS and single-cell cloned into 96-well plates. Clonal lines were expanded and genotyped by Sanger sequencing across the LRRC58 Cys319 locus, and the intended edit and absence of unintended indels were confirmed by amplicon deep sequencing analyzed with CRISPResso2[53]. The functional behavior of these cell lines was further confirmed by constitutive degradation of CDO1 in the presence of high concentrations of cystine as assessed by western blotting (**Fig. 5A**) as well as the CDO1 post-translational stability reporter as assessed by flow cytometry (**fig. S9A**).

**Cystine-depletion and repletion media.** Cystine-depletion medium was prepared using Dulbecco's modified Eagle's medium containing glucose and L-glutamine but lacking cysteine and methionine (US Biological, D9813). The medium was supplemented with sodium bicarbonate at 1.5 g liter$^{-1}$, L-methionine (Sigma-Aldrich, M5308) at 15 mg liter$^{-1}$, 10% dialyzed FBS (Thermo Fisher, 26400044), and 1% penicillin–streptomycin. L-cystine dihydrochloride (Sigma-Aldrich, C6727) was added at the indicated final concentrations. Before treatment, cells maintained in complete EMEM were washed with phosphate-buffered saline and transferred to cystine-depletion medium or to the same base medium supplemented with the indicated concentration of cystine. Cells were cultured under these conditions for the durations specified in the text and figure legends.

**Measurement of CDO1 post-translational stability.** Hep G2 cells expressing the CDO1–eGFP-IRES-mCherry post-translational stability reporter were exposed to the indicated experimental conditions. Cells were detached with 0.25% trypsin–EDTA, washed, and resuspended in PBS with 10% FBS containing 0.5 μg/mL DAPI immediately before analysis. Flow cytometry was performed using an LSRFortessa flow cytometer (BD Biosciences) equipped with either a high-throughput sampler or standard acquisition tubes. Data were analyzed using FlowJo versions 10 and 11. Cells were gated sequentially to identify intact cells, singlets, DAPI-negative viable cells, and mCherry$^{+}$ reporter-expressing cells, as shown in **fig. S5B**. CDO1 post-translational stability was quantified within the live, singlet, mCherry$^{+}$ population as either the eGFP:mCherry mean-fluorescence-intensity ratio and the mode-normalized eGFP fluorescence, as indicated in the figure legends.

**Recombinant protein expression and purification.** Baculovirus for protein expression of pAC8-derived constructs was generated by transfection of expression plasmids into Sf9 (Gibco) cells at a density of $0.9 \times 10^6$ cells ml$^{-1}$ in ESF 921 medium (Expression Systems). Viral titers were increased by three rounds of infection in Sf9 cells. StrepII–LRRC58 or StrepII-LRRC58(C319), CUL5, ELO-B, ELO-C and FLAG–RBX2 were co-expressed, while StrepII-Avi-CDO1 or StrepII-Avi-CDO1 (H147A) was expressed separately. Hi-5 cells were infected at a density of 1.8–$2.0 \times 10^6$ cells ml$^{-1}$ in SF-4 Baculo Express ICM medium (BioConcept) and collected 40–48 h post-infection by centrifugation (1,500 rpm, 20 min). The collected cells were resuspended in lysis buffer (50 mM HEPES/NaOH pH 7.4, 200 mM NaCl, 0.1% (v/v) Triton X-100, 1 mM TCEP) supplemented with protease inhibitors and lysed by sonication, and the supernatant was treated with benzonase for 10 min. The cell lysate was clarified by ultracentrifugation (40,000 rpm, 1 h) and filtered. Filtered lysate was applied to Strep-Tactin XT 4Flow resin (IBA) equilibrated in lysis buffer. The resin was washed with wash buffer (50 mM HEPES/NaOH pH 7.4, 500 mM NaCl, 1 mM TCEP) and bound protein was eluted with elution buffer (50 mM HEPES/NaOH pH 7.4, 200 mM NaCl, 50 mM biotin, 1 mM TCEP). Further cleanup was done with anion exchange chromatography (buffer A: 50 mM HEPES/NaOH pH 7.4, 1 mM TCEP, buffer B: 50 mM HEPES/NaOH pH 7.4, 750 mM NaCl, 1 mM TCEP) followed by SEC (SEC buffer: 50 mM HEPES/NaOH pH 7.4, 150 mM NaCl, 1 mM TCEP). FLAG–eGFP–LRRC58 and FLAG–eGFP–LRRC58(C319A) were each co-expressed with ELOB and ELOC in Hi-5 cells (Gibco), while FLAG-CDO1 was expressed separately. FLAG-tagged proteins were purified as described above with lysis and wash buffer instead containing 200 mM NaCl and no TCEP. Lysate was then applied to anti-FLAG resin (Genscript) and eluted (50 mM HEPES/NaOH pH 7.4, 200 mM NaCl, 150 μg ml$^{-1}$ FLAG peptide) prior to anion exchange and SEC.

**Co-immunoprecipitation binding assays with recombinant protein.** Preassembled recombinant FLAG-tagged CRL5$^{LRRC58}$ (2 μg) and recombinant Strep-CDO1 (0.2 μg) were combined in 1 mL binding buffer (25 mM Tris-HCl pH 7.5, 150 mM NaCl , 5% glycerol, 1% NP-40, ± EDTA or TPEN) with protease inhibitors, supplemented with cysteine, cystine, or other additives to constitute the indicated conditions, and preincubated for 1 h at 4 °C with end-over-end rotation. 30 μL samples were collected and saved following 1-hour preincubation as Input/Load. Anti-FLAG M2 magnetic beads were then added and samples were incubated for a further 1 h at 4 °C with end-over-end rotation. Beads were washed 3x in binding buffer, and bound protein was eluted with 5x bead volume of 150 ng/μL 3X FLAG peptide. Samples were analyzed by western blot.

**Western blotting.** Samples were isolated using radioimmunoprecipitation (RIPA) lysis buffer (Sigma R0278) or Pierce IP lysis buffer (Pierce Cat #87787) with protease inhibitors (Complete Mini, EDTA-free Protease Inhibitor Cocktail, Roche, 11836170001). Homogenates were centrifuged at 21,000g for 10 min at 4 °C, and the supernatants were used for subsequent analyses. Protein concentration was determined using the Pierce BCA assay. Protein lysates were denatured in NuPAGE 4x LDS Sample buffer (Invitrogen NP0007), resolved by 4–12% NuPAGE Bis-Tris SDS–PAGE (Invitrogen), and transferred to a polyvinylidene difluoride (PVDF) membrane using an iBlot 2 or 3 (Invitrogen).

Primary antibodies — anti-CDO1 (12589-1-AP, Proteintech, 1:1,000); anti-CUL5 (Bethyl Laboratories, A302-173A, 1:500); anti-ELOB (Proteintech, 10779-1-AP, 1:500); anti-ELOC (Proteintech, 12450-1-AP, 1:1,000); anti-vinculin (Cell Signaling Technologies, 4650 or 18799, 1:1,000); anti-Strep (Abcam, ab76949, 1:4000); anti-tubulin (Cell Signaling Technologies, 11H10, 2125, 1:1000); and anti-FLAG M2 (F1804, Sigma-Aldrich, 1:1,000) — were diluted in Tris-buffered saline containing 0.05% Tween-20 (TBS-T) and 5% BSA. Membranes were incubated with primary antibodies overnight at 4 °C. HRP-conjugated secondary antibodies (Promega anti-mouse W402B and anti-rabbit W401B, 1:5,000) were diluted in TBS-T containing 5% BSA. Signal was visualized with enhanced chemiluminescence (ECL) substrate (Thermo Scientific SuperSignal West Pico PLUS, 34580). Densitometry was performed in ImageJ 1.53.

**TR-FRET assay to measure affinity**

eGFP-LRRC58-ELOB-ELOC (WT), eGFP-LRRC58(C319A)-ELOB-ELOC, and biotinylated CDO1 were prepared as previously described[23]. To measure the $K_d$ value of complex formation, 200 nM biotinylated CDO1 and 2 nM of Tb-coupled streptavidin were mixed in an assay buffer with 25 mM HEPES, 150 mM NaCl, 1 mM fresh TCEP, 0.5% BSA, and 0.05% Tween-20 (mixture 1). Serial dilutions of eGFP–LRRC58–ELOB–ELOC (WT-LRRC58 or C319A-LRRC58) at a final concentration ranging from 9.8 nM to 20 μM were prepared in the same buffer (mixture 2). A total of 7.5 μl each of mixtures 1 and 2 were added to each well in a 384-well microplate and incubated for 1 h at room temperature in the dark and read on a Pherastar FS (BMG) plate reader with a 337/490/520 nm filter set. Background fluorescence due to eGFP–LRRC58–ELOB–ELOC was calculated by repeating the assay without CDO1 and subtracting the signal for the corresponding eGFP–LRRC58–ELOB–ELOC concentration. The ratio of emissions at 520/490 from five cycles were averaged as the final TR-FRET reading. $K_d$ values were calculated using one-site specific binding and plotted in GraphPad Prism 10.

**TR-FRET with cystine**

eGFP-LRRC58-ELOB-ELOC (WT), eGFP-LRRC58(C319A)-ELOB-ELOC, and biotinylated CDO1 were prepared as previously described[23]. Purified WT and mutant protein were buffer-exchanged into assay buffer (25 mM Tris-HCl pH 7.4, 150 mM NaCl, ± 1 mM EDTA, 0.5% BSA, 0.05% Tween-20) containing L-cystine at twice the final assay concentration (0, 125, 250, or 500 μM) and incubated on ice. Biotinylated CDO1 (200 nM) was buffer-exchanged into cystine-free assay buffer, combined with Tb-streptavidin (2 nM), and split ± 2 mM DTT. Equal volumes of LRRC58 and $CDO1_{Tb}$ were combined to a final concentration of 2265 nM LRRC58-ELOB-ELOC, 0–250 μM L-cystine, and ± 1 mM DTT, incubated 1 h at room temperature in the dark, and read on a Pherastar FS (BMG). The 520/490 nm emission ratio, averaged over five cycles, was normalized to the 0 μM cystine condition per genotype and analyzed in GraphPad Prism 10 and 11.

**Cryo-EM sample preparation and data collection**

Purified $CRL5^{LRRC58}$ (22.87 μM) was mixed with 34.31 μM CDO1 (a 1:1.5 molar ratio) in a final volume of 500 μl in SEC buffer (50 mM HEPES/NaOH pH 7.4, 150 mM NaCl, 1 mM TCEP). The mixture was incubated on ice for 1 h, and then the assembled complex was purified from unbound protein by SEC. Quantifoil R 1.2/1.3 Cu 300 grids were discharged at 20 mA for 1 minute (EMS GloQube) and loaded on a plunge freezer (Leica EMGP2) at 10 °C and 90% humidity. For the CDO1-LRRC58 sample, the pre-formed complex was mixed with β-OG detergent for a final concentration of 20 μM protein and 12.5 mM β-OG in a volume of 4 μL. The sample was applied to the grid, and after a 10 second pre-blot time was blotted for 5s and then plunged into liquid ethane. The Apo-LRRC58 sample was prepared in the same way. For the cystine-LRRC58 sample, $CRL5^{LRRC58}$ was buffer exchanged into 25 mM Tris-HCL pH 7.4, 150 mM NaCl, 250 μM L-cystine using a Zeba spin desalting column. Following a 2-hour incubation on ice, the cryo-EM sample was prepared as described above. All datasets were collected on a 300 kV FEI Titan Krios microscope located at the HHMI Janelia Research Campus. Movie collection was performed with a total exposure dose of 50 e-/Å2 and exposure time of 3.52463s. The CDO1-LRRC58 dataset was collected over 7 days, with half the collection time having the grid tilted to 30°. The Apo-LRRC58 dataset was collected over 3 days, and the Cystine-LRRC58 dataset was collected over two sessions totaling 6 days.

**Cryo-EM image processing**

All dataset processing and model-building was done using SBGrid-supported software cryoSPARC (v 4.5 and 5.0), Coot (v 0.9.8.92), and Phenix (v 2.2.6143)[54]. Detailed overviews of the processing workflows of all datasets are also described in **fig. S2, S4**, and **S6**.

**Model Building and deposition**

The CDO1-$CRL5^{LRRC58}$ composite map was generated using the Phenix Combine Focus Maps functionality from local refined maps separately masked around CDO1-LRRC58-ELOB-ELOC and CUL5-RBX2. This volume was then used for subsequent model building. An initial model was generated using AlphaFold3 and a cryo-EM model of CUL5-RBX2 (PDB: 6V9I), and the model was rigid-body fit into the cryo-EM density using UCSF ChimeraX. Manual adjustment of the model was done in Coot in conjunction with several rounds of real-space refinement in Phenix. The final CDO1 maps (composite, CDO1-LRRC58-ELOB-ELOC local, CUL5-RBX2 local, consensus) were deposited (EMD-79119, EMD-79114, EMD-79115, EMD-79116) alongside the model (PDB: 38SU). The final Apo-$CRL5^{LRRC58}$ (EMD-79124) and Cystine-$CRL5^{LRRC58}$ (EMD-79125) maps were deposited separately.

**Cryo-EM map comparisons and RMSD and angle analysis**

All cryo-EM maps were first fit into the composite CDO1-$CRL5^{LRRC58}$ map to globally align. The LRR domain (residues 11-247) of LRRC58 was first rigid-body fit into the Cystine-$CRL5^{LRRC58}$ map separately from the LRRC58 BC-box domain and the CUL5-ELOB-ELOC-RBX2, which were able to be fit as a single unit. RMSD and rotational angle values were calculated for each separately in ChimeraX (v1.9). Structural figures were also prepared in UCSF ChimeraX.

**Hydrogen Deuterium Exchange-Mass Spectrometry (HDX-MS) experiments**

CRL5–LRRC58 and CDO1 constructs were prepared as 20 μM stock solutions for HDX-MS analysis. Deuterium labeling was initiated with a 20-fold dilution (19 μL) in a $D_2O$ labeling buffer (25 mM Tris, ± 1 mM EDTA, 150mM NaCl, pD 7.0, 99 % $D_2O$) starting from a 20 μM protein solution (1 μL). After each labeling time (2 seconds, 5 seconds, 10 seconds, 1 minute, 10 minutes) at 20 °C, the labeling reaction was quenched by the addition of 20 μL of ice-cold quenching buffer (2 M guanidinium chloride, 0.8 % (v/v) formic acid, pH 2.3, $H_2O$) and analyzed immediately. For cystine-bound complexes, a super-saturated cystine solution (~460 μM) was preincubated with the protein for ~2 h prior to labeling, and the labeling buffer likewise contained ~460 μM cystine. For the CRL5–LRRC58(or CRL5–$LRRC58^{C319A}$)–CDO1 complexes, proteins were combined at a 1:1 molar ratio before labeling.

LC/MS: Deuterated and control samples were digested online at 15 °C using an AffiPro Nepenthesin II column (AffiPro, AP-PC-004). The cooling chamber of the UPLC system, which housed all the chromatographic elements, was held at 0.0 ± 0.1 °C for the entire time of the measurements. Peptides were trapped and desalted on a VanGuard Pre-Column trap [2.1 mm × 5 mm, ACQUITY UPLC BEH C18, 1.7 μm (Waters, 186002346)] for 3 minutes at 100 μL/min. Peptides were then eluted from the trap using a 5% –35% gradient of acetonitrile over 10 minutes at a flow rate of 100 μL/min, and separated using an ACQUITY UPLC HSS T3, 1.8 μm, 1.0 mm × 50 mm column (Waters, 186003535). The back pressure averaged ~12,950 psi at 0 °C and 5% acetonitrile 95% water, 0.1% formic acid. The error of determining the deuterium levels was ± 0.5 Da in this experimental setup. To eliminate peptide carryover, a wash solution (1.5 M guanidinium chloride, 0.8% formic acid and 4% acetonitrile) was injected over the Enzymate column during each analytical run. Mass spectra were acquired using a Waters SELECT SERIES Cyclic IMS QTOF instrument coupled to an I-class nano ACQUITY UPLC system and a HDX manager. The mass spectrometer was calibrated with direct infusion of a solution of major mix at flow rate 10 μL/ min (Waters, 186008113). Leucine enkephalin (Waters, 186006013) at 250 femtomole/ μL at a flow rate of 5 μL/ min was used as lock mass for post-acquisition mass accuracy correction. Spectra were acquired between 50 to 2000 m/z with capillary voltage 3.0 kV, trap collision energy at 4 V, sampling cone at 40 V, transfer CE ramping from 15 to 50 V, source temperature of 80 °C and desolvation temperature of 450 °C. For cIMS, a single-pass IMS with 10 ms injection, 3 ms separation, and 34 ms ejection /acquire with a sequence was used. The peptides were identified using undeuterated control samples using PLGS 3.0.3 (Waters Corporation). Peptide masses were identified from searches using non-specific cleavage of a custom database containing the sequence of LRRC58, LRRC58 C319A variant, CDO1, CUL5, Elongin B and C, and RBX2. Raw MS data were imported into DynamX 3.1.1 (Waters Corporation) and filtered with minimum consecutive products of 2, minimum number of products per amino acid of 0.25. Those peptides meeting the filtering criteria were further processed automatically by DynamX followed by manual inspection of all spectra and data processing. The relative amount of deuterium in each peptide was determined by subtracting the centroid mass of the undeuterated form of each peptide from the deuterated form, at each time point, for each condition. These deuterium uptake values are used to generate all uptake graphs and difference maps and are shown in **data S1**. The error of determining the average deuterium incorporation for each peptide was at or below ± 0.5 Da. Deuterium levels were not corrected for back exchange and thus reported as relative[55].

**HDX-MS data visualization**
Differential HDX-MS data are presented as the per-peptide difference in deuterium uptake ($\Delta D$, Da) between two states, plotted against residue number at a series of labeling times. Positive $\Delta D$ indicates deprotection (increased exchange) and negative $\Delta D$ indicates protection (reduced exchange) in the experimental state relative to the reference state. A significance threshold was defined as the peptide-averaged measurement error (typically ±0.5 Da); differences within this threshold are not considered significant. Deuterium levels were not corrected for back-exchange and are reported as relative values. For structural mapping, $\Delta D$ values at a single representative timepoint (specified in each figure legend) were projected onto the corresponding cryo-EM structural model. Only peptides with $|\Delta D|$ exceeding the significance threshold were considered significant. For each such peptide, the N-terminal residue (n−1) was excluded because its backbone amide undergoes rapid back-exchange and does not reliably report uptake, as were proline residues, which lack an exchangeable amide. Each remaining residue was assigned the significant peptide of largest $|\Delta D|$ covering it and colored on a binned diverging scale of five steps per direction (blue, protection; red, deprotection); residues that were covered but below the significance threshold are shown in white, and residues lacking peptide coverage or corresponding to prolines are shown in gray. Because colors are assigned per peptide, the maps reflect peptide-level, not residue-level, resolution. To emphasize the magnitude of significant differences, the polypeptide backbone was rendered as a variable-radius tube (worm representation) in which the radius scaled continuously with $|\Delta D|$ across the significant range; residues below the significance threshold and those lacking data were rendered at a uniform minimum radius. Color bin edges and the corresponding radius mapping were set per dataset according to the dynamic range of $\Delta D$ observed and are reported in the corresponding figure legend; magnitudes should therefore be compared within, but not across, panels.

**Immunoprecipitation of FLAG-LRRC58.** LRRC58 knockout (KO) Hep G2 cells expressing FLAG-tagged WT or C319A LRRC58 were cultured in cystine-depleted medium overnight, pretreated with 10 μM MG132 for 1 h, and then treated with 10 μM MG132 plus cystine for the indicated duration. Cells were lysed in IP Lysis buffer (25 mM Tris-HCl pH 7.4, 150 mM NaCl, 1% NP-40, 5% glycerol, ±1 mM EDTA) with protease inhibitors, and protein concentration was determined by BCA assay. FLAG-LRRC58 was immunoprecipitated using anti-FLAG M2 magnetic beads and eluted with 3X FLAG peptide.

**Quantification of LRRC58 cysteine reversible covalent modification.** Following overnight cystine depletion, cells were treated with 10 μM MG132 plus 200 μM cystine for 0, 2, or 3 h (MG132 present for a total of 4 h in all conditions). FLAG-LRRC58 immunoprecipitation was performed as described above. Each eluate was split into two aliquots: one was directly alkylated with iodoacetamide (35 mM final) to cap free thiols, while the other was first reduced with TCEP (6.25 mM final). Both were cleaned up by single-pot, solid-phase-enhanced sample preparation (SP3 ref. [56]) bead capture to remove excess TCEP and alkylated with IAM (35 mM final). Following a second round of SP3 bead capture, samples were digested on-bead with trypsin/LysC. The resulting peptides were desalted and analyzed by parallel reaction monitoring (PRM) on a timsTOF HT (shown in **table S4, fig. S5H**). Reversible oxidation of each cysteine-containing peptide was calculated as 100% × [1 − (A_−TCEP/A_+TCEP)], where A_+TCEP and A_-TCEP denote the integrated peak areas of the iodoacetamide-labeled peptide in matched aliquots analyzed with or without TCEP reduction, respectively.

**LRRC58 cysteine zinc occupancy by differential TPEN chelation.** LRRC58 KO Hep G2 cells expressing WT FLAG-LRRC58 were treated with 10 μM MG132 plus 200 μM cystine for 0 or 2 h (MG132 present for a total of 3 h in all conditions), followed by FLAG-LRRC58 immunoprecipitation as described above. Each eluate was split into two aliquots and alkylated with iodoacetamide (IAM, 10 mM final) in the presence or absence of the zinc chelator TPEN (1 mM final), to compare cysteine reactivity with and without zinc chelation. Samples were incubated for 15 min at 37°C followed by 45 min at room temperature in the dark, then processed by SP3 bead cleanup, digested with trypsin/LysC, desalted, and analyzed by PRM on a timsTOF HT (**table S4** and **fig. S5H**). Increased IAM labeling in the +TPEN condition relative to control identifies cysteines whose reactivity is normally restricted by zinc coordination. Percent zinc occupancy was calculated as 100% × [1 − (A_−TPEN/A_+TPEN)] where A_+TPEN and A_-TPEN denote the integrated peak areas of the iodoacetamide-labeled peptide in matched aliquots with or without TPEN chelation, respectively. Percent zinc occupancy was bounded from 0-100% for display. C301/C308/C309 were resolved as one tryptic peptide. C331 and C335 were not detected.

**Hep G2 Mass spectrometry acquisition and proteomics data analysis.** Desalted peptides from the cysteine-reactivity, zinc-occupancy, and K111[GG] proteomics workflows were separated on an Evosep One (Evosep) using the 30 sample-per day method with a PepSep C18 Column (15 cm × 150 μm, particle size 1.5 μm) and analyzed on a timsTOF HT (Bruker). Untargeted assays used DDA and data were analyzed in FragPipe (v23, MSFragger) using a custom database including UniProt human plus FLAG-LRRC58 and CDO1-eGFP. . Targeted assays used prm-PASEF monitoring the peptides listed in **table S4**. Peptides were quantified in MS2 by integrating the listed transition, either manually in Compass DataAnalysis 6.2 or in Skyline (26.1.0.057).

**Quantification of LRRC58 K111[GG]**. Following overnight cystine depletion, cells were pre-treated with 10 μM MG132 for one hour followed by the addition of 200 μM cystine with 10 μM MG132 for the indicated time. FLAG-LRRC58 immunoprecipitation was performed as described above. Eluates were alkylated with iodoacetamide (IAM, 14 mM final), cleaned up by SP3 bead cleanup, and digested with trypsin/LysC. They were then desalted and analyzed by parallel reaction monitoring (PRM) on a timsTOF HT (Bruker, **table S4**). The LGGPSALPK[+114.0429]GLAQSPLC[+57.0215]R peptide (K111[GG]) was identified by the presence of multiple transitions (**fig. S8A,B**). K111[GG] peptide intensity was quantified from its most intense product ion, $y11^{2+}$ (m/z 670.86), of the triply charged precursor at m/z 646.0192 and integrated manually in Compass DataAnalysis 6.2 or in Skyline (26.1.0.057). K111[GG] intensity was normalized to total LRRC58 protein abundance via normalization to LRRC58 peptides DLTYDPPTLLELAAR and LSVSTETLESELEAR.

**Quantification of diglycine modification at other LRRC58 sites.** Samples were prepared as described above, analyzed by data-dependent acquisition (DDA) and processed in FragPipe v23 with variable modifications including GlyGly-Lys (+114.04293 Da). Ubiquitylation is reported as MaxLFQ intensity from the combined K[114.0429]site analysis, normalized to the average across conditions for that site. K99[GG] and K314[GG] were only detected in a minority of samples in some experiments and were therefore not quantified.

**Hep G2 cell metabolite extraction and sample preparation.** Confluent Hep G2 cells in 6-well plates were cultured for 12 h in cystine-depleted medium or medium containing 200 µM $^{13}C_6$,$^{15}N_2$-L-cystine. All subsequent steps were performed on ice. The media was aspirated, and cells were washed twice with ice-cold PBS, with any residual liquid removed. Ice-cold 90% methanol (300 µL per well) was added, and cells were scraped into the solvent and transferred to microcentrifuge tubes. Extracts were vortexed, precipitated at −80 °C overnight, vortexed again, and clarified by centrifugation at 17,000g for 15 min at 4 °C. The supernatant was transferred to autosampler vials and analyzed on an Orbitrap Exploris 120 (Thermo Fisher Scientific) using amide column chromatography in positive ion mode. For targeted quantification of free cysteine, an aliquot of the same extract was reduced with TCEP (0.5 mM final, 1 h at room temperature) and alkylated with N-ethylmaleimide (NEM; 0.5 mM final, 1 h at room temperature) to cap free thiols as the cysteine–NEM adduct, and analyzed on a T3 column in positive ion mode. See "**Metabolite LC-MS acquisition and data analysis"** section for details on chromatography and mass spectrometry.

***C. elegans*** **metabolite extraction and sample preparation.** All steps were performed on ice or at 4 °C unless otherwise noted. Frozen worm pellets were resuspended in 250 µL ice-cold 100% methanol by vortexing and pipetting, sonicated in a cold room using a Diagenode Bioruptor UCD-200 Sonicator (high setting, 0.5 s on/0.5 s off, 10 min), rotated end-over-end for 1 h at 4 °C, and precipitated at −80 °C overnight. Extracts were clarified by centrifugation at 17,000g for 15 min at 4 °C, and the supernatant was collected and dried by vacuum centrifugation. Dried metabolites were resuspended in ice-cold 95% methanol, clarified again at 17,000g for 15 min at 4 °C, and supernatant transferred to autosampler vials for LC-MS analysis. For targeted metabolite profiling, samples were analyzed on an Orbitrap Exploris 120 (Thermo Fisher Scientific) using amide column chromatography in positive and negative ion modes. For targeted quantification of free cysteine, an aliquot of the same extract was reduced with TCEP (0.5 mM final, 1 h at room temperature) and then alkylated with N-ethylmaleimide (NEM; 0.5 mM final, 1 h at room temperature) to cap free thiols as the cysteine–NEM adduct, and analyzed on a T3 column in positive ion mode. See "**Metabolite LC-MS acquisition and data analysis."** section for details on chromatography and mass spectrometry.

**Metabolite LC-MS acquisition and data analysis.** LC-MS analyses of metabolite extracts were performed on a Vanquish HPLC System coupled to an Exploris 120 mass spectrometer equipped with a HESI ion source (Thermo Fisher Scientific). The LC system was controlled by Chromeleon software (Thermo Fisher Scientific), and the MS was controlled by Xcalibur software (Thermo Fisher Scientific). Samples were analyzed using two chromatography methods described as follows. When using Waters ACQUITY UPLC HSS T3 column (150 x 2.1 mm, particle size 1.8 µm) kept at 35 °C with a flow rate of 0.4 mL/min, 3 min isocratic mobile phase A (water with 0.1% formic acid) followed by a 14.5-min linear gradient from 1% to 99% mobile phase A – mobile phase B (acetonitrile with 0.1% formic acid) was used. When separating using Waters XBridge Premier BEH Amide column (100 mm × 2.1 mm, particle size 2.5 µm), solvent A was 95% acetonitrile/ 5% water with 10 mM ammonium acetate and 0.1% formic acid, and solvent B was 50% acetonitrile/ 50% water with 10 mM ammonium acetate and 0.1% acetic acid. A flow rate of 0.4 mL/min was used and A/B gradient was as follows: isocratic at 5% B for 1.5 min, linearly increasing to 98% B at 9 min, 98% B for 1 min, shifting back to 5% B in 0.1 min and holding at 5% B until 15 min. Mass spectrometer parameters: spray voltage positive/negative switching electrospray ionization (ESI) mode +3.5 kV or negative ESI -2.5 kV; ion transfer tube temperature 320 °C; vaporizer temperature 350 °C; sheath gas 50 AU; auxiliary gas 10 AU; sweep gas 1 AU; RF Lens 70%; resolution 120,000; AGC target standard. The instrument was calibrated with FlexMix calibration solution (Thermo Fisher Scientific). m/z range was 70−800. MS1-MS2 was acquired for a 'pool' sample that was generated by equally mixing all samples: for MS1, orbitrap was operated at a resolution of 60,000; for MS2, top 4 ions at an isolation window of 1.2 m/z with 10 s exclusion (± 5 ppm) were triggered for MS2, orbitrap resolution was 30,000, and the following normalized collision energy was applied 20%, 40%, 60%.

The resulting files were analyzed by Compound Discoverer (Thermo Fisher Scientific, version 3.3) using the default metabolomics workflow including online database search (mzCloud, Mass List, and ChemSpider). Compounds with a minimum peak shape score of 3 in 60% of sample files were included for analyses. Quantification of LC-MS data was performed through peak detection and integration functions in Freestyle software (Thermo Fisher Scientific, version 1.8 SP2 QF1) using mass ranges of calculated [M-H]- or [M+H]+ m/z ± 5 ppm. In the case of quantifying stable isotope-incorporated metabolites, m/z window was manually examined to ensure exclusion of natural isotopes. In the case of *C. elegans* metabolite data, targeted metabolite abundances were normalized to the median signal intensity of all named metabolites derived from the paired untargeted metabolomics analysis of the same sample to correct for differences in worm number between samples.

**Sequence alignment of LRRC58 and LRR-2.** Full-length protein sequences of human LRRC58 (UniProt Q96CX6, residues 1–371) and *C. elegans* LRR-2 (UniProt Q9N4Z5, residues 1–375) were aligned with MAFFT v7.525[57] via the EMBL-EBI Job Dispatcher[58] using the BLOSUM62 scoring matrix, a gap-opening penalty of 1.53, a gap-extension penalty of 0.123, and two rounds of iterative refinement (--maxiterate 2, --retree 2). The resulting alignment was rendered with ESPript 3.2[59], with similarity shaded by %Equivalent (global score 0.7, difference score 0.5) and residues colored by physicochemical property. Conserved cysteines and lysine and the corresponding residue positions were annotated manually.

**AlphaFold modelling.** Full-length UniProt sequences for *C. elegans* CUL-5, RBX-2, ELB-1, ELC-1, LRR-2 and CDO-1 were submitted as a single six-chain job to the AlphaFold Server (AlphaFold 3, Google DeepMind[60]). The top-ranked of five output models was used without refinement (seed 903587327). For **Fig. 5I**, the LRR-2 and CDO-1 chains were superposed onto the corresponding chains of the human $CRL5^{LRRC58}$–CDO1 cryo-EM structure in UCSF ChimeraX 1.11.1 (matchmaker, iterative pruning at 2.0 Å). Owing to interdomain flexibility at the BC-box linker, the LRR solenoid and C-terminal domain were superposed independently (split at LRR-2 residue 257); the two fits are related by a rotation of 37.9°. The dashed line in **Fig. 5I** indicates the unmodelled linker.

***C. elegans* and bacterial maintenance and strains.** *C. elegans* strains used: N2 Bristol (wild-type), obtained from the *Caenorhabditis* Genetics Center (University of Minnesota); FGC125 (*Y42G9A.3*(syb11237[C330A])) and FGC132 (*Y42G9A.3*(syb11238[C330A]), generated via CRISPR Cas9 by Sunnybiotech and backcrossed three times to N2. For the lifespan tests, both strains were used. For the remaining assays the FGC125 strain was used. Wild-type *E. coli* strain BW25113 was obtained from the *E. coli* Genetic Resource Center. Bacteria were grown at 37 °C in LB medium ± kanamycin (50 μg/mL) at 200 rpm for 16 hours. Worms were maintained on Nematode Growth Media (NGM) agar plates at 20 °C, seeded with 120 μL of an *E. coli* BW25113 overnight culture three days before worms were added.

***C. elegans* development assay.** Worms were synchronized by pooling gravid *C. elegans* worms in M9, followed by a bleach:NaOH (7:8, v/v) treatment, to free the eggs from the adult worms. Eggs were washed four times with fresh M9 after bleaching and hatched overnight in fresh M9 at room temperature on a shaker. Approximately 100 larval stage one (L1) worms were added to the experimental plates, and developmental stages were determined 48 hours later.

***C. elegans* fecundity assay.** Single worms were transferred to experimental plates at larval stage four (L4). Worms were transferred to fresh experimental plates every 24 hours, and offspring were counted on each plate 48 hours after the worm was added.

***C. elegans* lifespan assay.** Per condition, synchronized L4 worms were transferred to three experimental plates, 30 worms per plate. To prevent the development of eggs, 5-fluoro-2'-deoxyuridine (ThermoScientific, L16497.ME) was added to the plates to a final concentration of 1 μM. Worms were manually scored every two days and were transferred to fresh plates on day four of adulthood. Worms were censored if they went missing or showed vulval protrusion or bagging.

**Collection of worms for metabolomics and proteomics.** Per condition, ~2000 synchronized worms were transferred at the L4 stage to experimental plates and allowed to grow until day 1 of adulthood. Worms were washed off plates with M9 and washed three additional times with fresh M9 to remove bacteria, before pellets were flash frozen in liquid nitrogen.

***C. elegans* proteomics.** Protein was recovered from the pellets post-metabolite extraction. Pellets were washed with 1 mL ice-cold 100% methanol and centrifuged at 17,000g for 5 min at 4 °C; all liquid was removed and pellets were air-dried for 10 min. Pellets were solubilized in 250 μL 4% SDS, 25 mM Tris-HCl pH 7.5 by pipetting, and 750 μL benzonase buffer (50 mM HEPES pH 8, 150 mM NaCl, 1 mM $MgCl_2$, Benzonase Nuclease at 1 μL/mL) was added (final 1% SDS), followed by incubation for 10 min at room temperature. Protein concentration was determined by BCA assay. For each sample, 10 μg protein in 45 μL 1% SDS (pH ~8) was reduced with 5 mM DTT for 30 min at 37 °C and alkylated with 14 mM iodoacetamide for 45 min at room temperature in the dark. 2 μL SP3 beads were added, followed by 50 μL 100% ethanol, and samples were incubated for 5 min at room temperature with vortexing to bind protein to the beads. Beads were immobilized on a magnetic rack, the supernatant was discarded, and beads were washed three times with 80% ethanol. Protein was digested on-bead with 0.5 μg trypsin and 0.3 μg LysC in 15 μL ammonium bicarbonate buffer overnight at 37 °C. Beads were immobilized on a magnet, and the peptide-containing supernatant was transferred to a fresh tube, acidified with 100 μL 1% formic acid, and desalted by StageTip.

Desalted peptides samples were analyzed using an Evosep One (EVOSEP) liquid chromatography system coupled to a timsTOF HT (Bruker) mass spectrometer. Samples were reconstituted in water containing 0.1% formic acid and loaded onto Evotip. Peptides were separated using a PepSep C18 Column (15 cm × 150 μm, particle size 1.5 μm) and a 30 SPD method (44-min gradient) on Evosep

One. CaptiveSpray source was operated at 1600 V with dry gas flow 3 L/min at 180 °C. The mass spectrometer was operated under data-independent acquisition with parallel accumulation serial fragmentation (DIA-PASEF) mode. MS parameters: polarity positive, scan m/z range 100–1700, mobility (1/K0) range 0.60–1.60 V·s/cm2, ramp time 75 ms, accumulation time 75 ms. MS/MS was performed using optimized window settings with 1.7 s cycle time estimate, 1 MS1 ramp, 20 MS/MS ramps, 60 MS/MS windows, 350.7–1250.6 Da mass range, and 0.6–1.45 V·s/$cm^2$ mobility range (1/$K_0$). Linear mobility-dependent collision energy was applied with 20 eV at 0.60 V·s/$cm^2$ and 59 eV at 1.6 V·s/$cm^2$.

Data analyses were performed using DIA-NN (version 2.3.0) using developer-recommended settings (default except using 15 ppm mass errors). In silico predicted library was generated from a FASTA file retrieved from Uniprot (www.uniprot.org, UP000001940). Protein groups with valid quantities in ≥ 75% of replicates in at least one genotype were retained; missing values were imputed within each genotype to the median of detected values, or to an instrument floor (0.9 × the lowest observed intensity) when a genotype was undetected. Intensities were $\log_2$-transformed and compared by two-sided Welch's t-test with Benjamini–Hochberg FDR correction; proteins with $|\log_2$ fold-change$| > 1$ and FDR < 0.05 were considered differentially abundant.

**Statistical analysis.**

Analyses were performed in GraphPad Prism 10 and 11. Statistical tests are stated in the figure legends. $P < 0.05$ was considered significant.

**References (continued from Main Text):**


48 Vakulskas, C. A. *et al.* A high-fidelity Cas9 mutant delivered as a ribonucleoprotein complex enables efficient gene editing in human hematopoietic stem and progenitor cells. *Nat Med* **24**, 1216-1224 (2018). https://doi.org/10.1038/s41591-018-0137-0

49 Zuris, J. A. *et al.* Cationic lipid-mediated delivery of proteins enables efficient protein-based genome editing in vitro and in vivo. *Nat Biotechnol* **33**, 73-80 (2015). https://doi.org/10.1038/nbt.3081

50 Doman, J. L., Sousa, A. A., Randolph, P. B., Chen, P. J. & Liu, D. R. Designing and executing prime editing experiments in mammalian cells. *Nat Protoc* **17**, 2431-2468 (2022). https://doi.org/10.1038/s41596-022-00724-4

51 Anzalone, A. V. *et al.* Search-and-replace genome editing without double-strand breaks or donor DNA. *Nature* **576**, 149-157 (2019). https://doi.org/10.1038/s41586-019-1711-4

52 Chen, P. J. *et al.* Enhanced prime editing systems by manipulating cellular determinants of editing outcomes. *Cell* **184**, 5635-5652 e5629 (2021). https://doi.org/10.1016/j.cell.2021.09.018

53 Clement, K. *et al.* CRISPResso2 provides accurate and rapid genome editing sequence analysis. *Nat Biotechnol* **37**, 224-226 (2019). https://doi.org/10.1038/s41587-019-0032-3

54 Herre, C. et al. Introduction of the Capsules environment to support further growth of the SBGrid structural biology software collection. Acta Crystallogr D Struct Biol **80**, 439-450 (2024). https://doi.org/10.1107/S2059798324004881

55 Wales, T. E. & Engen, J. R. Hydrogen exchange mass spectrometry for the analysis of protein dynamics. *Mass Spectrom Rev* **25**, 158-170 (2006). https://doi.org/10.1002/mas.20064

56 Hughes, C. S. *et al.* Single-pot, solid-phase-enhanced sample preparation for proteomics experiments. *Nat Protoc* **14**, 68-85 (2019). https://doi.org/10.1038/s41596-018-0082-x

57 Katoh, K. & Standley, D. M. MAFFT multiple sequence alignment software version 7: improvements in performance and usability. *Mol Biol Evol* **30**, 772-780 (2013). https://doi.org/10.1093/molbev/mst010

58 Madeira, F. *et al.* The EMBL-EBI Job Dispatcher sequence analysis tools framework in 2024. *Nucleic Acids Res* **52**, W521-W525 (2024). https://doi.org/10.1093/nar/gkae241

59 Robert, X. & Gouet, P. Deciphering key features in protein structures with the new ENDscript server. *Nucleic Acids Res* **42**, W320-324 (2014). https://doi.org/10.1093/nar/gku316

60 Abramson, J., Adler, J., Dunger, J. et al. Accurate structure prediction of biomolecular interactions with AlphaFold 3. *Nature* **630**, 493–500 (2024). https://doi.org/10.1038/s41586-024-07487-w

**Table S1. (separate file)**

Processing and refinement statistics for CDO1-CRL5$^{LRRC58}$, Apo-CRL5$^{LRRC58}$ and cystine treated CRL5$^{LRRC58}$.

**Table S2. (separate file)**

Proteins significantly altered in comparative proteomics between LRR-2(C330A) and WT *C. elegans*.

**Table S3. (separate file)**

Oligonucleotides and DNA sequences used in this study.

**Table S4. (separate file)**

Details of parallel reaction monitoring used in this study.

**Movie S1.**

Movie displaying the motion of the LRR arm of LRRC58 in the Apo vs cystine treated CRL5$^{LRRC58}$ structures.

**Data S1. (separate file)**

This file contains the tabulated data plotted in the figures.
